# A neural circuit for context-dependent multimodal signaling in *Drosophila*

**DOI:** 10.1101/2024.12.04.625245

**Authors:** Elsa Steinfath, Afshin Khalili, Melanie Stenger, Bjarne L. Schultze, Sarath Nair Ravindran, Kimia Alizadeh, Jan Clemens

## Abstract

Many animals, including humans, produce multimodal displays by combining acoustic with visual or vibratory signals [1–4]. However, the neural circuits that coordinate the production of multiple signals in a context-dependent manner are unknown. Multimodal behaviors could be produced by parallel circuits that independently integrate the external cues that trigger each signal. We find that multimodal signals in *Drosophila* are driven by a single circuit that integrates external sensory cues with internal motivational state and circuit dynamics. *Drosophila* males produce air-borne song and substrate-borne vibration during courtship and previous studies have identified neurons that drive courtship and singing, but the contexts and circuits that drive vibrations and coordinate multimodal signaling were not known [5–11]. We show that males produce song and vibration in distinct, largely non-overlapping contexts and that brain neurons that drive song also drive vibrations with cell-type specific dynamics and via separate pre-motor pathways. This circuit also coordinates multimodal signaling with ongoing behavior, namely locomotion, to drive vibrations only when the male’s vibrations can reach the female. A shared circuit facilitates the control of signal dynamics by external cues and motivational state through shared mechanisms like recurrence and mutual inhibition. A proof-of-concept circuit model shows that these motifs are sufficient to explain the behavioral dynamics. Our work shows how simple motifs can be combined in a single neural circuit to select and coordinate multiple behaviors.

---

Social communication is inherently multimodal. During conversations, we are not mere loud-speakers that emit speech but coordinate our words with dynamical facial expressions and other body gestures. Gestures produced in congruence with speech rhythms can improve comprehension [12, 13] whereas reducing multimodality, as in phone calls, can impair it [14, 15]. Multimodal communication is not unique to humans [1, 2] but also prevalent in other animals. For instance, monkeys [3], birds [16], frogs [17], or grasshoppers [18] combine acoustic signals with visual displays [3, 19, 20], while many insects combine sound with substrate-borne vibrations [4, 21–25]. Effective multimodal communication requires the production of the appropriate sequence or combination of signals contingent upon the context, for example, coordinating movements with a dance partner [26, 27].

Due to the multifaceted nature of multimodal signaling, the underlying brain circuits have mainly been studied by isolating single components of this behavior [3, 6, 28–32], but their contribution to the coordination of multimodal signals is not well understood. Moreover, the mechanisms by which these circuits integrate external cues for context-appropriate signaling [8, 33] and coordinate signaling with ongoing behaviors such as respiration and locomotion is poorly understood [34, 35]. At one extreme, parallel circuits could independently integrate the specific external cues required to trigger different behaviors [1]. Alternatively, a single integrated circuit could trigger multiple behaviors and signal coordination arises from the interaction between external sensory inputs, internal motivational state, and circuit dynamics [10, 36, 37].

Here, we address the issue of multimodal signaling in *Drosophila melanogaster*. During courtship, male flies chase females while producing both air-borne song and substrate-borne vibration to attract their attention [5, 38]. Song is produced by extending and fluttering one wing resulting in two distinct modes: a sine song characterized by sustained sinusoidal oscillations with a frequency around 150 Hz, and a pulse song consisting of trains of short pulses with two distinct shapes, produced at a regular interval of around 40 ms [39]. Substrate-borne vibrations are associated with abdominal quivering and are pulsatile like the pulse song, but with a longer interval of 150–200 ms [5]. Both signals are evaluated by the female and influence her mating behavior [40, 41]. However, how the male brain coordinates air-borne song and substrate-borne vibration is unknown.

In the *Drosophila* brain, sexual behaviors are controlled by sexually-dimorphic neurons that express the transcription factors *fruitless* or *doublesex* [5, 42–45]. The neural circuitry underlying courtship song production is well understood with central neuron types P1a and pC2l integrating social cues—chemical, visual, acoustic—to drive persistent courtship and singing [10, 30, 36, 46, 47] in the ventral nerve chord (VNC) via at least two descending neurons (DNs), pIP10 [6] and pMP2 [7]. The choice between the two song modes is driven by the relative activity of these DNs and by circuit dynamics in the VNC [10, 11].

In contrast, the behavioral contexts and neural circuits that drive vibration in *Drosophila* males are unknown. It is unclear to what extent song and vibration are produced simultaneously or sequentially since recordings of both signals with sufficient temporal resolution in naturally interacting animals are lacking. Because vibrations are associated with abdominal quivering rather than wing movements like the song [5, 48] they are likely generated by a separate motor program.

## Simultaneous recordings of song and vibration during courtship in *Drosophila*

To assess the coordination of song and vibration, we designed a behavioral chamber that can reliably record song and vibration simultaneously (Fig. 1A–C, S1C, modified from [8, 49]). Microphones tiling the behavioral setup floor were covered by a thin paper serving as a substrate for the flies to walk on and for transmitting both signal types. We discriminated song and vibration pulses based on their interval differences whereby song pulses arrive at intervals between 30 and 45 ms, and inter-vibration intervals (IVIs) are much longer and range between 140 and 180 ms (Fig. 1D). Using laser vibrometry, we observed IVIs matching previous readouts of vibrations ([5], Fig. S1A, B). By recording high-resolution video of courtship in a smaller chamber and analyzing the movement of the abdomen during vibrations using SLEAP pose tracking ([50], Fig. S1D, E) we confirmed that the vibration pulses are associated with the previously reported abdominal quivering [5].

**Figure 1:**
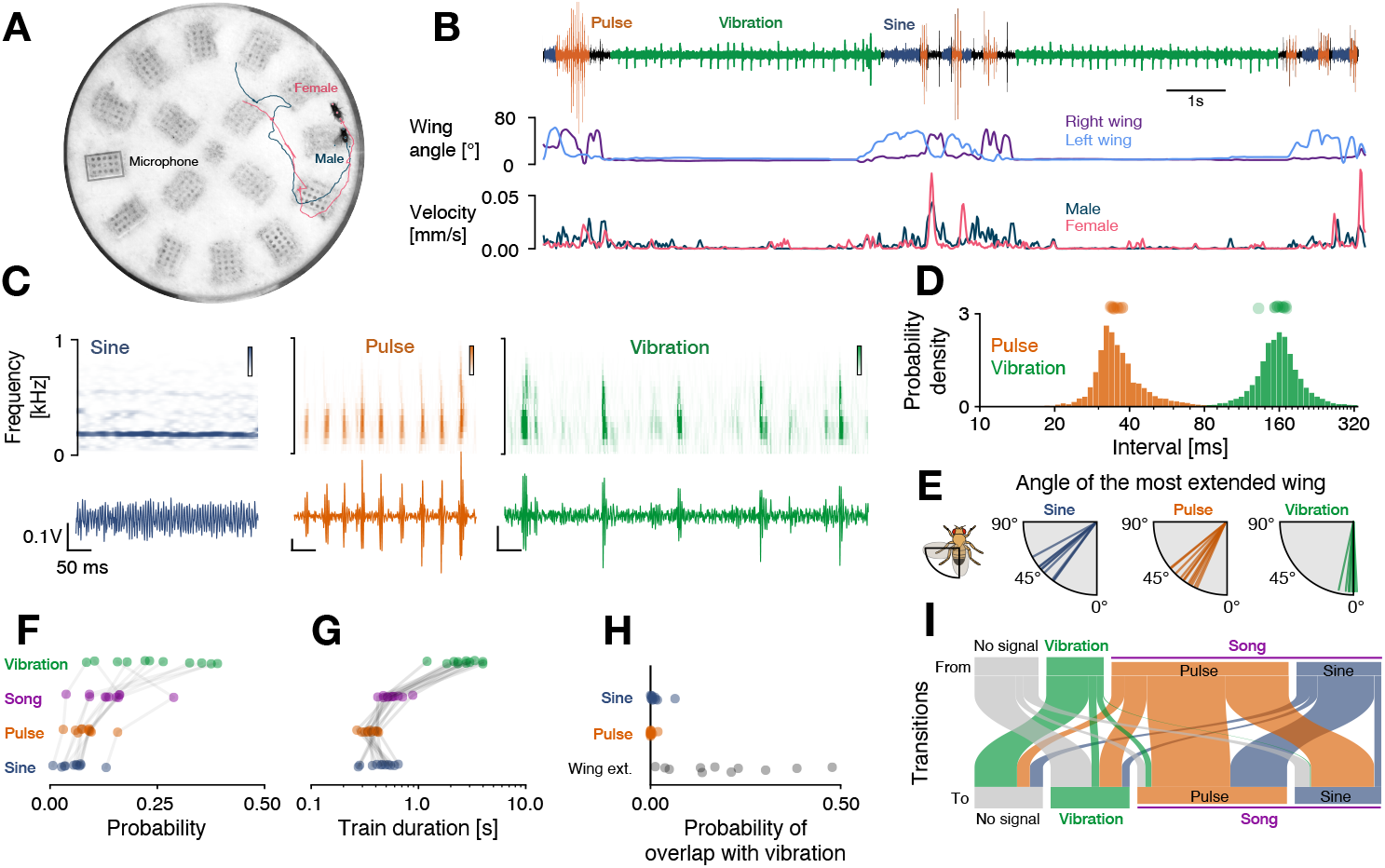
Drosophila males produce two multimodal signals—song and vibration—during courtship. **A** Behavioral chamber with a male (blue) courting a female fly (pink) and tracked poses (dots) and walking trajectories (lines). One of the 16 microphones embedded in the floor is marked with a grey box. **B** Audio trace (top) from one of the microphones with sine song (blue), pulse song (orange), and vibrations (green) alongside behavioral cues extracted from pose tracking: The angle of the male’s left and right wing (middle) as well as male and female velocity (bottom). **C** Waveforms (bottom) and spectrograms (top) for sine song (blue), pulse song (orange), and vibrations (green). **D** Distribution of intervals between song pulses (orange, N=27310) and between vibrations (green, N=16785). Dots on top show median values for each male. Intervals between song pulses (35.5±11.4 ms, median±interquartile range (IQR)) are much shorter than intervals between vibrations (160±41 ms). **E** Median angle of the most extended wing during sine (58±8°, median±IQR), pulse (48±9°) and vibration (12±5°). Values close to 0° correspond to no wing extension. Males rarely extend their wing during vibrations. **F** Probability of producing sine (6±3%, mean±standard deviation), pulse (8±3%), song (14±6%), vibration (24±10%), and no signal (62±11%) during courtship. Males produce more vibrations than song (p=0.02). **G** Duration of sine songs (460±145 ms), pulse trains (355±79 ms), song bouts (562±129 ms), and vibration trains (2785±944). Vibration trains are longer than song bouts (p=5 ·10^−4^) **H** Overlap between vibrations and sine song (0.012±0.017), pulse song (0.002±0.006) orwing extensions (0.19±0.14). **I** Transitions between no signals (grey), vibration (green), pulse (orange), and sine (blue). Line width is proportional to the probability of transitioning from one signal (top) to another (bottom). Transitions between the song modes (pulse and sine) are more frequent than between song and vibrations p=5 10^−4^. N=11 males in D–I. All reported p-values from one-sided Wilcoxon tests. Reported summary statistics correspond to mean±standard deviation (std.) unless noted otherwise.

### Male flies dynamically switch between song and vibration during courtship

With access to song and vibration produced by the male during courtship, we next characterized the coordination between these two signals. During courtship, males vibrated twice as much compared to singing, and the vibration bouts were longer than song bouts (Fig. 1F, G). Song is produced using uni-lateral wing extensions while vibrations do not require the wings (Fig. 1E, S1G, H). Although 19% of vibrations occurred while the wing was extended, males rarely sang and vibrated at the same time (1%) (Fig. 1H, S1F) indicating that the male is physically able to simultaneously sing and vibrate but chooses not to overlap both signals.

The male switched dynamically and non-randomly between sine, pulse and vibrations (Fig. 1I). Transitions between the song modes (sine, pulse) were more frequent (26% of all transitions), than transitions between song and vibration (only 7% of all transitions). Moreover, while pulse and sine were sequenced into bouts with no or very short pauses, vibrations were separated from song by a pause of around 1 second (Fig. S2). This temporal coordination of song and vibration suggests that these two signals are produced in distinct behavioral contexts. To identify these contexts, we next linked recordings of song and vibration with video tracking of the courtship interactions using computational modeling.

### Locomotion and distance of the female fly determine signal choice

The choice between sine and pulse song is based on female feedback [8, 9, 39] and our analyses of the transitions between song and vibration suggest that this might also be true for vibrations (Fig. 1I). To identify the cues that inform the male’s choice between song and vibration, we employed generalized linear models (GLMs) using the dynamics of social cues extracted from the male and female tracking data to predict the male’s choice between song, vibration, or no signal (Fig. 2A, B).

**Figure 2:**
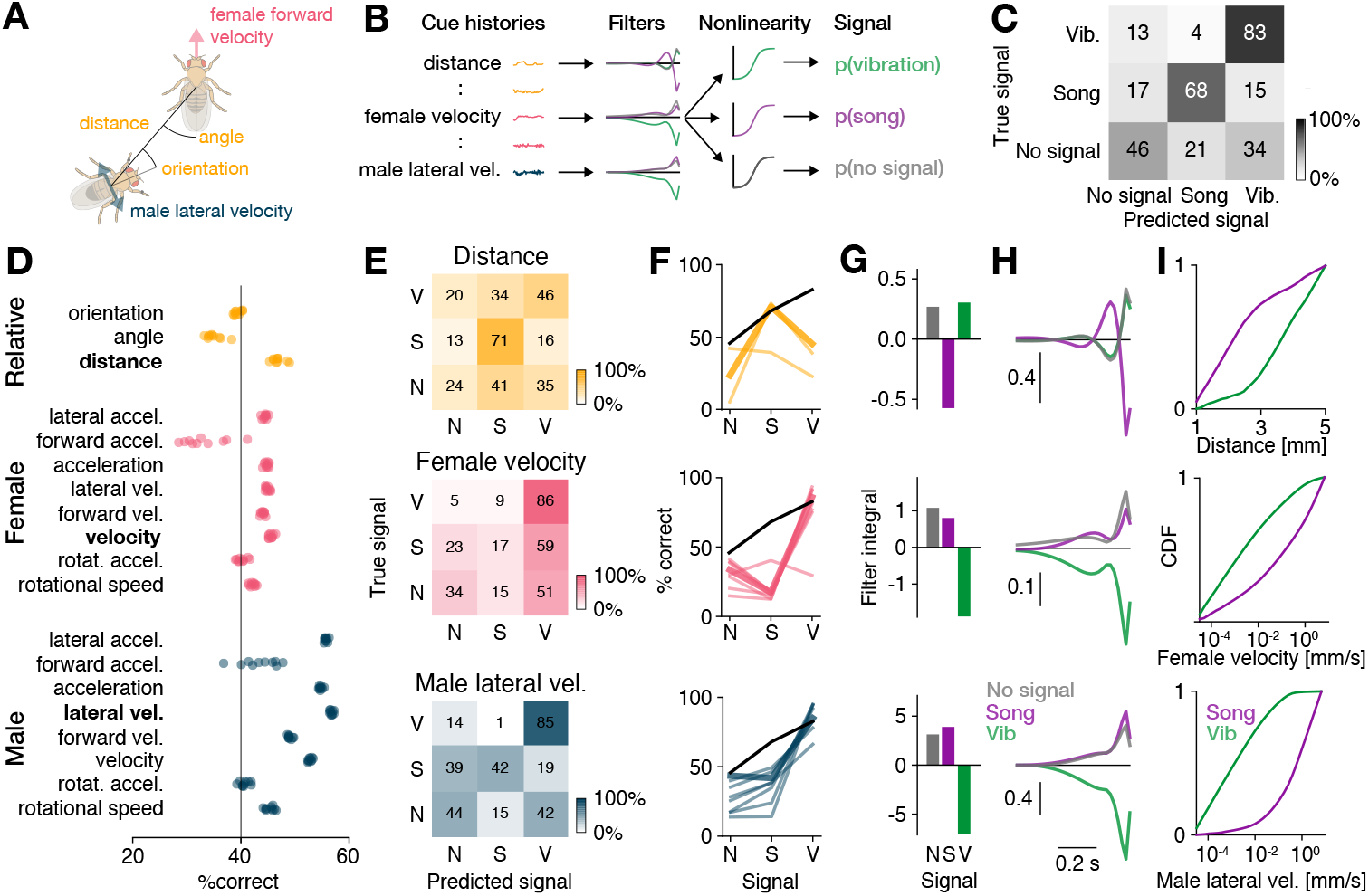
Locomotion and distance predict signal choice. **A** Examples of feedback cues used to predict the male’s signal choice. **B** Signal choice (song, vibration, no signal) was predicted using the cues histories (A) from one second preceding each time point. Choice relevant temporal cue patterns were detected using filters, with one filter per cue and signal type. The filtered cues are then passed through a nonlinearity that yields the probability of observing each signal. **C** Confusion matrix for a model fitted to predict the male’s signal choice from all cues. Shading and numbers indicate the classification percentage (see color bar). **D** Predictive performance (% correct)of individual male (blue), female (pink), and relative (yellow) cues. Dots correspond to result from 10 model fits from independent train-test splits. **E** Confusion matrices for the prediction of signal choice (N - no signal, S - song, V - vibration) for the most predictive male cue (lateral velocity, bottom), female cue (female velocity, middle), and relative cue (distance, top). Shading and numbers indicate the classification percentage (see color bar). **F** Signal-wise performance for male (bottom), female (middle), and relative (top) cues. Male cues predict vibrations very well and song moderately. Female cues only predict vibrations well and relative distance predicts song well. Thick colored lines correspond to the best cue for each cue group shown in E. Black lines show the performance of the multi-feature model from C. See also Fig. S3A. **G** Integral over the filters for each signal for the cues shown in E. Small male (bottom) and female velocity (middle) values predict vibration. Small male-female distances (top) predict song. **H** Filter shapes of the cues shown in E. The distance filter for song changes its sign from positive to negative, indicating that a reduction in distance drives song. **I** Cumulative density functions (CDFs) for the cues shown in E. Vibrations are produced at low velocities (bottom, middle) and song is produced at smaller distances (top).

With only rare confusions between song and vibration we were able to determine that feedback cues determine the choice between song and vibration (Fig. 2C). To assess the contribution of individual cues to the signal choice, we fitted individual models for each cue (Fig. 2D–F, S3A, B) and found that models fitted with male or female locomotor cues predicted vibrations best, with 83-92% accuracy, while relative cues like distance and orientation were less predictive (<50%). In contrast, song was predicted best by the relative cues distance and orientation (71%), less well by male cues (38-49%), and poorly by female cues (12-18%). These findings indicate that male and female movement patterns are the strongest predictors of vibration production during courtship.

We then determined how the cues influence signal choice by examining the integral of each cue’s filter. If the sign of the integral is positive, then high cue values (e.g. large distances) promote the signal; if the sign is negative, then the cue suppresses the signal. The filters for the best male and female predictors—female velocity and male lateral velocity—were positive for song and no signal but negative for vibrations (Fig. 2G, H). This trend was consistent for all locomotion filters (Fig. S3B) indicating that males tend to vibrate when they or the female are slow or stationary, and they tend to sing when either the male or female are moving (Fig. 2I, S3B–D). The observed association between stationarity and male vibration production is not due to limitations in our recording setup (Fig. S1E) and is consistent with previous findings linking female immobility to increased male vibration behavior [5].

The filter for distance, the cue most predictive of singing, was negative for singing and positive for vibrations, indicating that males vibrate when farther away from the female and sing when in closer proximity (Fig. 2G–I, S3C–F). In addition, the distance filter for song changed its sign from positive to negative, indicating that a reduction in distance to the female drives singing (Fig. 2H). This is consistent with singing frequently preceding copulation attempts, during which a previously stationary male moves closer to the female [51]. Distance is known to determine the choice between song types [8, 39], as well as the amplitude of song [52]. It also determined the choice between song and vibration, indicating its centrality for courtship signal choice. Interestingly, the context in which males vibrate—slow and far from the female—was previously interpreted as a disengaged state [9]. Having access to vibrations during courtship, we found that part of this ‘passive’ state is not idle, but that the male actively signals to the female.

### Stationarity is necessary and sufficient to drive vibrations in males

The statistical models of male signal choice showed that stationarity predicts vibrations (Fig. 2). However, it is possible that other behaviors that females primarily perform when stationary (e.g. grooming) could be the cause for vibrations. We therefore causally tested the role of stationarity by manipulating locomotion during courtship. According to the behavioral models, stopping the male or the female should increase the probability of observing vibrations, while inducing locomotion should suppress vibrations (Fig. 2G–I). To not interfere with the male’s signaling ability, we optogenetically manipulated female walking behavior during courtship.

We first stopped the female by expressing GtACR1, an inhibitory channelrhodopsin, in all motor neurons (using the vGlut driver) [53]. Stopping the female increased vibrations by 30% (Fig. 3A, B). Conversely, inducing female walking by optogenetically activating the DNp28 neurons [54, 55] nearly abolished vibrations (Fig. 3C, D). These causal interventions therefore confirmed that stationarity is necessary and sufficient for vibrations. Further, singing was best predicted by male-female distance (Fig. 2F), but distance changed only little when stopping the female (Fig. S4B). Distance did increase when inducing female locomotion and this weakly suppressed singing (Fig. S4C), demonstrating that controlling female locomotion only weakly affected singing behavior (Fig. S4A–C) consistent with the behavioral models (Fig. 2E, F). In summary, locomotion controls vibrations.

**Figure 3:**
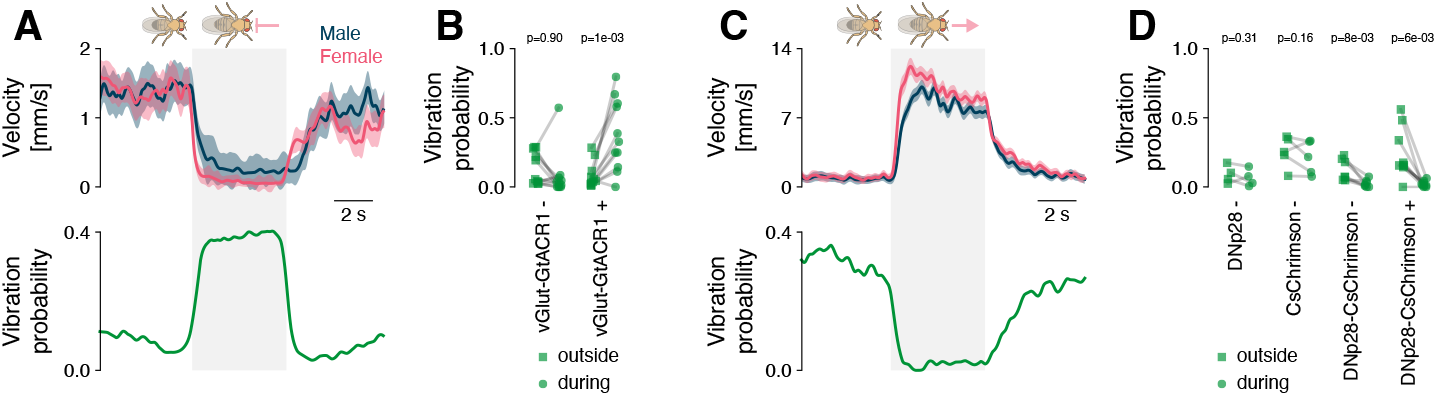
Female immobility is a necessary and sufficient trigger for male vibrations. **A** Optogenetic inactivation (grey) of all motor neurons (MNs) in a female courted by a wild type male stops the pair (top, male/female velocity blue/pink) and triggers male vibrations (bottom). Females expressed GtACR1 in all glutamatergic neurons. Optogenetic stimulus 525 nm at 14 mW/cm^2^. **B** Average vibration probability outside of (squares) and during (circles) optogenetic inactivation of the MNs. Control females (vGlut-GtACR1-) had the same genotype but were not fed all-trans retinal, a co-factor required to make GtACR1 light sensitive. Lines connect data from the same pair during the different epochs (vGlut-GtACR1 atr-N=11, atr+ N=11). P-values from a paired Wilcoxon test of the hypothesis that the vibration probability increases due to female slowing. **C** Optogenetic activation (grey) of DNp28 neurons in a female courted by a wild type male accelerates the pair (top, male/female velocity blue/pink) and suppresses male vibrations (bottom). Optogenetic stimulus 625 nm at 89 mW/cm^2^. **D** Average vibration probability outside of (squares) and during (circles) optogenetic activation of DNp28. Lines connect data from the same pair during the different epochs (DNp28-Gal4+ N=4, UAS-Chrimson+ N=5, DNp28-Chrimson-N=7, DNp28-Chrimson+ N=9). P-values from a paired Wilcoxon test of the hypothesis that the vibration probability decreases due to female acceleration. Lines and shaded areas in A and C show the mean±standard error of the mean. A ‘+’/’-’ after the genotype names in B and D indicates the presence/absence of all-trans retinal in the food.

Although we genetically controlled female locomotion, the male chases the female and his movement is tightly correlated to her movement (Fig. 3A, C), in short, stopping the female during courtship also stops the male. This correlation also explains why both male and female locomotor cues predict vibrations (Fig. 2F, S3A). However, male signal choice is more strongly determined by his own than by the female’s stationarity (Fig. 2D, S4D): Male velocity distributions are clearly distinct when he sings versus vibrates, while female velocity distributions overlap considerably during song or vibration. It is therefore likely that the male’s locomotor state controls the choice between song and vibration, and is not influenced by the female movement. This co-regulation of locomotion and signaling likely evolved because walking can interfere with the transmission and perception of vibrations [41].

### Central “song” neurons drive vibration with complex dynamics

Having shown that locomotion regulates the switch to and from vibration, we next asked how this switch is implemented in the fly brain. While the neurons in the central brain that drive singing have been identified [6, 10], cell types that drive vibration are unknown. To test whether song and vibration are driven by distinct or overlapping central circuits, we examined whether key neurons of the song pathway also drive vibrations.

Several cell types that express the sex-determination genes *doublesex* and *fruitless* [42–44, 56–58] integrate social cues and drive singing in males. We focused on two brain-local neurons and two descending neurons that drive singing when activated. The pC2l neurons in the central brain, process auditory and visual cues and elicit robust singing via a direct connection to the descending pIP10 neurons [7, 10, 30, 39, 45, 59, 60]. The P1a neurons [6, 10, 39, 47, 60] process pheromones [46, 61, 62] and likely receive input from pC2l neurons [10]. P1a neurons induce a persistent arousal state that can drive courtship and singing, or aggression, [63, 64] on two timescales: on the order of up to ten seconds, via slowly decaying activity in P1a itself [62] and on the order of up to a minute via a recurrent neural network downstream of P1a [36]. P1a neuron activation alone tends to yield only little song since it drives song indirectly, via a disinhibitory circuit motif [10, 36, 39, 63]. The decision to sing, encoded in the activity of pC2l and P1a type neurons, is relayed to premotor circuits in the VNC via at least two descending neurons: pIP10 and pMP2 [6, 59]. pIP10 neurons receive inputs from the pC2l neurons but the central inputs to pMP2 or downstream targets of P1a are unknown.

Activation of all *doublesex* and *fruitless* neurons induces vibrations [5], but specific cell types— and hence circuits—that drive vibrations were not known. We optogenetically activated P1a [63], pC2l [45], pIP10 [6], and pMP2 [7] in solitary males with varying light intensities and examined the time spent producing each of the communication signals—vibrations, pulse, sine—during and between activations (Fig. 4B). The activation of the descending neurons pIP10 or pMP2 drove song but no vibrations. However, the two central brain neurons P1a and pC2l elicited both song and vibration. Among males with activated pC2l neurons, 8 out of 25 vibrated, and all 35 males with activated P1a neurons vibrated. This suggests that multimodal signal generation is orchestrated by a shared neural circuit capable of driving both signals. Consequently, descending neurons engage distinct motor circuits in the ventral nerve cord, dedicated to either song production or vibration.

**Figure 4:**
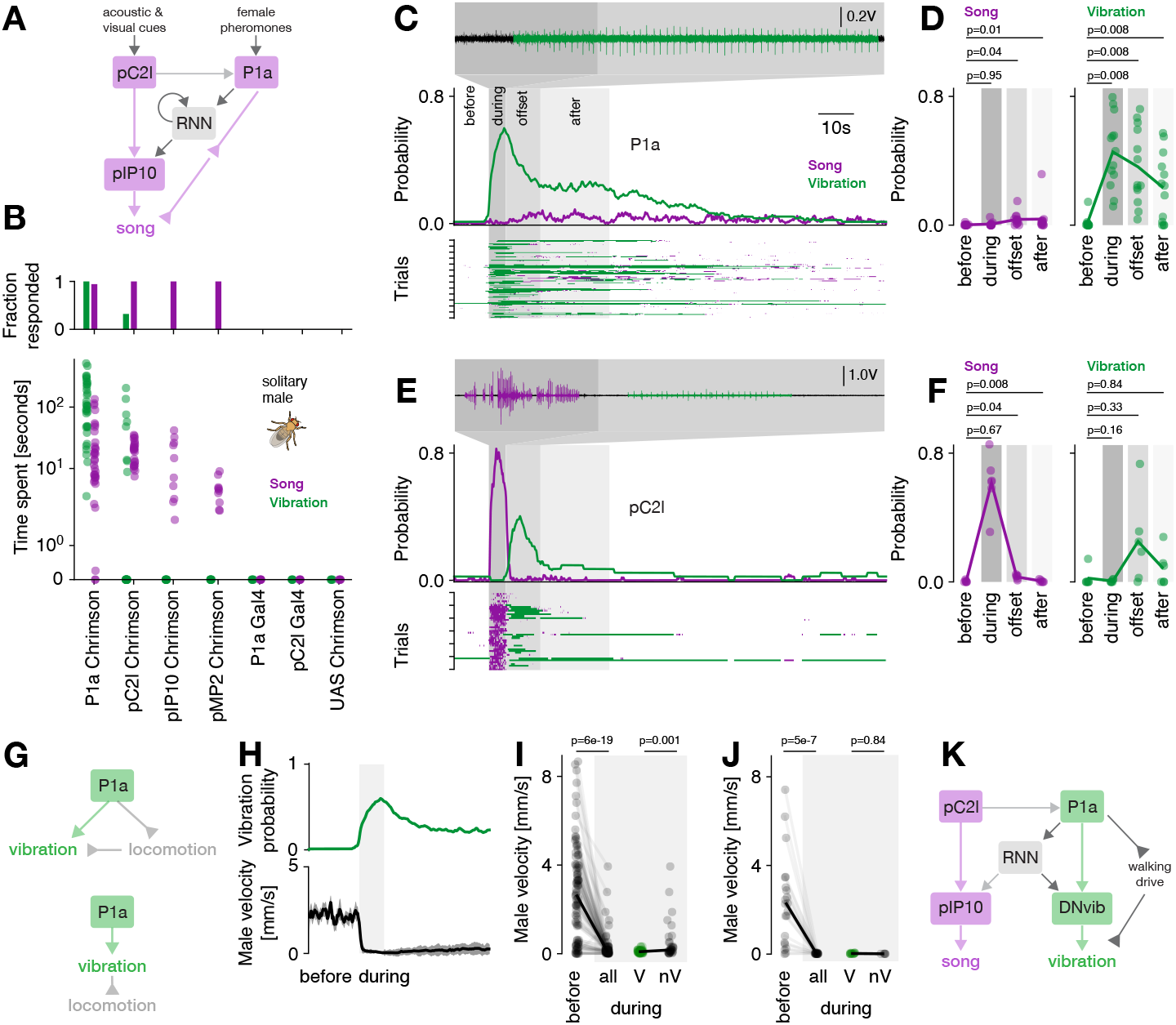
Dynamical multimodal signaling is controlled by a network that contains P1a and pC2l neurons. **A** The song circuit of *Drosophila*. The central neurons pC2l and P1a process social cues and trigger courtship and song. pC2l drives song via a connection to the descending neuron (DN) pIP10. Another DN with unknown inputs in the brain, pMP2, also drives song (not shown). P1a drives song indirectly, via a downstream recurrent neural network (RNN) and a disinhibitory circuit motif. Regular and inverted arrow heads indicate excitatory and inhibitory connections, respectively. **B** Song (purple) and vibration (green) evoked by optogenetic activation of P1a, pC2l, pIP10 and pMP2 across a range of light intensities. Bars (top) show the fraction of males that produced song (purple) or vibration (green) during an experiment. Dots (bottom) show the average time spent producing song (purple) or vibration (green) for individual males. Y-axis symlog scaled to include 0. N=35/25/10/10/6/5/5 males P1a/pC2l/pIP10/pMP2-Chrimson, three controls (P1a-Gal4, pC2l-Gal4, UAS-Chrimson). **C** Microphone recording (top), trial average probability (middle), and single trial raster (bottom) showing song (purple) and vibration (green) in response to optogenetic activation of P1a in solitary males (27 mW/cm^2^, N=13 flies, 7 trials/fly). Areas with different shades of grey delimit the different epochs analysed in D. **D** Probability of observing song (left) and vibration (right) in different epochs surrounding P1a activation (times relative to activation onset: before -10–0, during 0–5, offset 5–15, after 15–35 s) **E** Same as C but for optogenetic activation of pC2l (83 mW/cm^2^, N=6 flies, 7 trials/fly). **F** Same as D but for pC2l activation. **G** Two different hypotheses regarding the control of vibration and locomotion. Either, P1a independently controls vibration and suppresses locomotion (top). Or, P1a drives a single motor program that stops the male and makes him vibrate (bottom). **H** Vibration probability (top) and average male velocity (bottom) in response to optogenetic activation of P1a (same data as E). Nearly all males stop, but only 50% of the males vibrate. **I** Male velocity before and during optogenetic activation of P1a. Dots correspond to trials. Males are split into vibrating (green, V) and non-vibrating males (black, nV) based on whether they produced vibrations during the activation in that trial. **J** Same as I but for stronger P1a activation (209 mW/cm^2^, N=3). **K** Current working model of multimodal signaling in *Drosophila*. P1a drives vibrations directly and persistently, through direct and indirect (via RNN) connections with an unidentified descending neuron DNvib. In addition, P1a independently controls vibrations and locomotion to tie vibrations to phases of male stationarity. P-values in D, F from a Wilcoxon test testing the hypothesis that the probability of song or vibration increases. P-values in I, J from Mann-Whitney U tests of the hypothesis that P1a activation slows males, and that vibrating males are slower.

We next examined the dynamics with which P1a and pC2l drove multimodal signals. Activating P1a neurons [63] reliably induced vibrations that outlasted the activation for tens of seconds (Fig. 4C, D, Fig. S5A, B), independent of activation strength (Fig. S5E). Our sparse activation protocol also resulted in a few song bouts during and after activation. This implies that the persistent courtship state induced by P1a neuron activation jointly controls the multimodal courtship signals of song and vibration [36, 63]. By contrast, pC2l neuron activation reliably drove song (Fig. 4E, F, S5C, D). Interestingly, at the offset of strong activation, we observed vibrations lasting 5–10 s (Fig. 4F). pC2l neurons are known to produce sine song at activation offset [7, 10, 30] but this sine song is much shorter (<1 s) than the vibrations (5-10s) (Fig. S5D).

Thus, the “song circuit” comprised of P1a and pC2l neurons drove multi-modal signals. pC2l neurons directly drove song, P1a directly drove vibrations. The celltype-specific dynamics likely reflect differences in downstream connectivity. As pC2l neurons drive song via a direct connection to pIP10 [10, 65], we hypothesize that P1a neurons similarly drive vibrations via an unknown descending neuron (DNvib). Further, pC2l drives offset sine via its connection to P1a neurons, disinhibiting ventral nerve chord sine nodes [10]. We hypothesized that the offset vibrations are also driven through this pC2l-P1a connection and the DNvib.

### Central P1a neurons jointly control male locomotion and vibrations

Signaling needs to be coordinated with ongoing behaviors to ensure it’s efficacy, e.g, vocalizations are coordinated with breathing in vertebrates [34, 66]. Our behavioral analyses (Fig. 2, 3) showed that stationarity triggers vibrations, and P1a neuron activation is known to induce locomotor arrest in males [39, 63]. This suggests that P1a neurons not only drive multimodal signals but also coordinate them with locomotion. This could be attributed to P1a neurons either controlling locomotor state and vibrations in parallel or inducing a vibration motor program that inherently includes stopping the male (Fig. 4G). In the first case, P1a neuron activation should stop males, but not all stationary males should vibrate. In the other case, all males that stop upon P1a neuron activation should also vibrate. We therefore examined the association between P1a neuron activation, male locomotion, and vibrations. We find that P1a neuron activation induced locomotor arrest in solitary males [39] in almost all males (Fig. 4H, I). However, only 60% of the stationary males vibrated independent of activation strength (Fig. 4J), suggesting that P1a neurons do not induce a drive to vibrate which in turn stops males. Instead, P1a neuron activation induces two distinct motor programs: one that near-deterministically stops the male and puts him into “vibration mode” and another that then probabilistically triggers vibrations within this state. However, this does not rule out the possibility that locomotor state itself inhibits vibrations through an additional gating mechanism in moving males. Activation of pC2l neurons does not strongly affect locomotion, but males stop at activation offset, likely because pC2l neurons drive vibrations through P1a neurons (Fig. S5G). Thus, P1a neurons coordinate signaling with ongoing behavior—they stop males and induce vibrations.

## Mutual inhibition coordinates song and vibration

During natural courtship and during optogenetic activation, song and vibration rarely overlap (Fig. 1H), raising the question of how the song and vibration pathways interact downstream of P1a and pC2l neurons. A common circuit motif that prevents the simultaneous expression of two behaviors is mutual inhibition [67, 68] and might be at work downstream of P1a and pC2l. More specifically, we predicted that P1a neuron activation would suppress song since it drives vibrations, and pC2l neuron activation would suppress vibrations, given that it drives song (Fig. 5A, B). To unmask mutual inhibition between the song and vibration pathways, we activated P1a and pC2l neurons not in solitary males but in males paired with a female. We hypothesized that the presence of the female would drive P1a and pC2l neurons, consequently trigger courtship with song and vibration (Fig. 5C, D, S6A, B). Consistent with our prediction, P1a neuron activation strongly suppressed song (Fig. 5C, E) by interrupting singing in all flies, even in those that did not switch to vibrations (Fig. S6C). Conversely, pC2l neuron activation almost completely suppressed vibrations (Fig. 5D, F). Almost all flies that were vibrating in the five seconds prior to activation ceased their vibrations, even if they did not initiate singing behavior (Fig. S6D). These results show that mutual inhibition reduces the overlap between multimodal signals in *Drosophila*.

**Figure 5:**
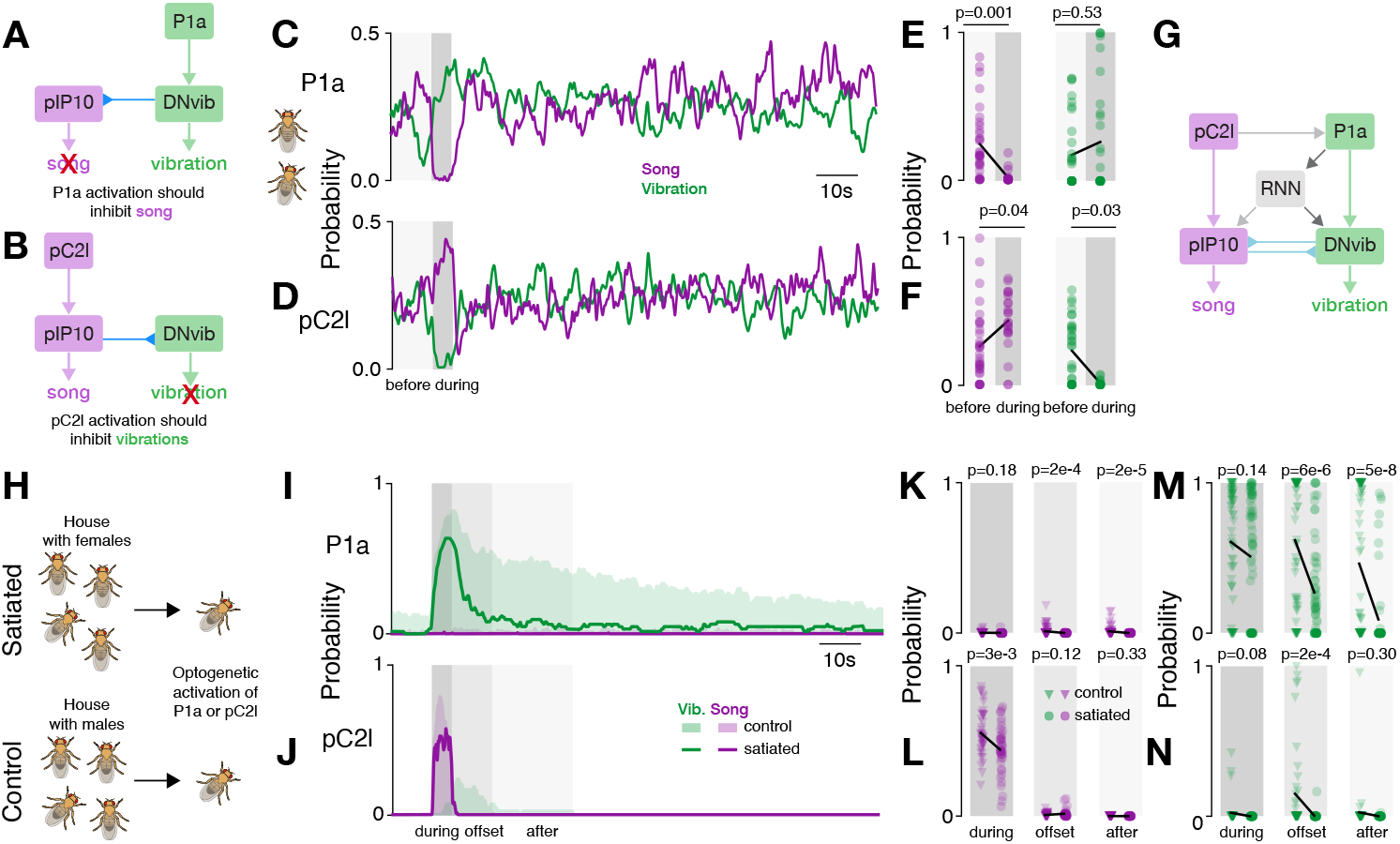
Coordination and modulation of song and vibration via mutual inhibition, female cues, and motivational state. **A, B** Hypothesized effects of mutual inhibition. Activation of P1a drives vibration and should inhibit song (A). Activation of pC2l drives song and should inhibit vibrations (B). For convenience, mutual inhibition is depicted as acting directly via the descending neurons, but it could also act downstream, in the ventral nerve cord. **C, D** Probability of song (purple) and vibration (green) in males courting a female during optogenetic activation of P1a (C) or pC2l (D). The presence of the female drives baseline signaling outside the activation window and unmasks the suppressive effect of mutual inhibition. P1a activation suppresses song and pC2l activation suppresses vibrations. Shaded areas indicate the time windows used for statics in E and F. For calculating the probabilities, only time steps during which the male courted the female were included. Light intensity 27 mW/cm^2^ at 625 nm. **E, F** Comparison of song (left) and vibration (right) in before (10 s) and during (5 s) activation of P1a (E) and pC2l (F) in males courting a female. P1a activation suppresses song and has no effect on vibrations in this context. Activation of pC2l increases singing and suppresses vibrations. The statistical tests only included trials in which the males courted the female in the windows before and during activation. P-values from two-sided Wilcoxon test. **G** Diagram of a working model of multi-modal signaling with mutual inhibition. **H** Males were sexually satiated by housing them with 10-15 virgins 4-6 h prior to the experiments. Control males were housed with 10-15 males. **I, J** Probability of observing song (purple) and vibration (green) in sexually satiated (lines) and naive (shaded areas) solitary males upon optogenetic activation of P1a (I) and pC2l (J). Gray shaded areas indicate time windows used for statics in K–N. **K, L** Comparison of song evoked in different time windows for P1a (K) and pC2l (L) in sexually satiated and control males. **M, N** Same as K, L but for vibrations. Data points in E, F and K–N correspond to trials and males. N males per genotype in F–F: 6 flies, G-I 4 flies, with 7 trials per male. Windows in E, F K–N were defined as follows: during (full 5 s of activation), offset (10 s after activation), after (10-30 s after activation). P-values in K–M from two-sided Mann-Whitney U tests. Black lines in E, F, K–M connect the medians between groups.

### Circuit dynamics bias signaling and can be overridden by female cues for context-appropriate signaling

Optogenetic activation engaged a circuit with strong autonomous dynamics (Fig. 4C–F): P1a neurons drive vibrations during and for tens of seconds after activation and only little song in solitary males. pC2l neurons drive a sequence of song during, and vibrations for 5–10 s after activation. However, signal dynamics during natural courtship with a female are much more variable (Fig. 1), because P1a and pC2l are activated by dynamical social cues from the female—P1a by contact and volatile pheromones [46, 61, 62] and pC2l by acoustic and visual cues [10, 30, 60]. For instance, the pulse to vibration transitions produced by pC2l activation (Fig. 4E) are rarely seen during natural courtship (Fig. 1I). To assess how dynamical social cues modulate the circuit’s autonomous dynamics during courtship, we assessed the data from activated P1a and pC2l neurons in males that courted a female (Fig. S6C, D). In the courting males, we found that activation of P1a or pC2l neurons did bias subsequent signaling towards vibrations. However, the bias was relatively weak and not as persistent as in solitary males (compare Fig. 4C, E). Thus, the circuit driving song and vibration in the central brain enables persistent yet flexible signaling. In the absence of social cues, activation of the circuit drives autonomous dynamics that enable persistent signaling. However, external cues can override these circuit dynamics to enable context-appropriate dynamical signaling.

### Song and vibration are under common motivational control

The persistence of courtship in *Drosophila* is driven by P1a neurons and modulated by sexual satiation, which reduces the initiation and persistence of courtship in males [69]. The effect of satiation is mediated by dopamine and leads to a reduced excitability of P1a neurons [62, 69] as well as less persistence in P1a neuron activity itself [62] and in the recurrent circuitry downstream of P1a neurons [36, 70]. One advantage of driving song and vibration through a shared circuit is that only a few circuit nodes need to be manipulated to globally up- or down-regulate multimodal signaling. However, direct effects of sexual satiation on singing and vibration have not been investigated. To assess whether motivational state modulates the persistence of both signals, we induced sexual satiation by allowing males to freely mate with females, and subsequently activated P1a and pC2l neurons (Fig. 5H). We found that sexual satiation strongly reduced the persistence of both song and vibration (Fig. 5I–N). Satiated males were less likely to vibrate after P1a neuron activation, and their tendency to sing was even further diminished (Fig. 5I, K, M). For pC2l activation, satiation weakly reduced the singing and almost completely abolished vibrations after activation offset (Fig. 5J, L, N). An effect of sexual motivation on P1a neurons has been demonstrated previously [62, 69, 70] and we now show that pC2l neurons were also subject to motivational control implying a global effect of motivation on the courtship circuit.

### A neural circuit model for multimodal signaling

Our experiments revealed a neural circuit that drives multimodal signals with complex and persistent dynamics. To test whether this circuit is indeed sufficient to explain the dynamics of multimodal signaling in *Drosophila*, we implemented a proof-of-concept circuit model (Fig. 6A, S7). The proposed model consisted of three major components: First, at the top of the hierarchy are pC2l and P1a neurons, which are activated by social cues (or optogenetically) and drive song and vibration (Fig. 4C, E). Direct connections between pC2l and P1a neurons and descending neurons mediated the immediate effects of social cues or optogenetic activation in our experiments. pC2l is directly connected to pIP10, which drives song in the VNC [65]. Given that P1a neurons strongly drove vibrations with little delay (Fig. 4C), we hypothesized that P1a neurons are connected to an unknown vibration descending neuron, that we called DNvib. Second, all indirect effects of optogenetic activation—the vibrations at the offset of pC2l neuron activation as well as the persistent song and vibration after P1a activation—were mediated by P1a neurons. P1a neurons are known to drive slow circuit dynamics in two ways: Intrinsically, through the slow decay of P1a neuron activity itself, which lasts 5-10 s [62]. And extrinsically, through a recurrent neural network (RNN) downstream of P1a neurons that maintains activity for several tens of seconds [36, 63]. The timescale of the intrinsic decay matched the timescale of offset vibrations after pC2l neuron activation. Behavioral [10] and female connectome data (Fig. S8) [71, 72] suggest that pC2l neurons likely weakly connect to P1a. Activation of pC2l would thus sufficiently drive P1a to induce the slowly decaying activity in P1a neurons, but not strongly enough to engage the RNN downstream of P1a neurons. Activation of the RNN requires strong and direct activation of P1a neurons and mediates the long-term persistence of multimodal signals via connections to the descending neurons for song and vibration. Lastly, the inhibitory cross-talk between song and vibration was mediated by mutual inhibition downstream of pC2l and P1a neurons, likely at the level of the descending pathways or in the premotor centers in the VNC [7]. In the model, we implemented mutual inhibition between pIP10 and DNvib neurons. Activation of pC2l neurons activates pIP10 neurons and pIP10 neurons drive song but also inhibit DNvib neurons and hence vibrations. Conversely, activation of P1a neurons activates DNvib neurons which drive vibrations and inhibit pIP10 neurons and thereby song. Adaptation and noise in the mutual inhibition can enable bistable dynamics [68], which in our model leads to switching between song and vibration.

**Figure 6:**
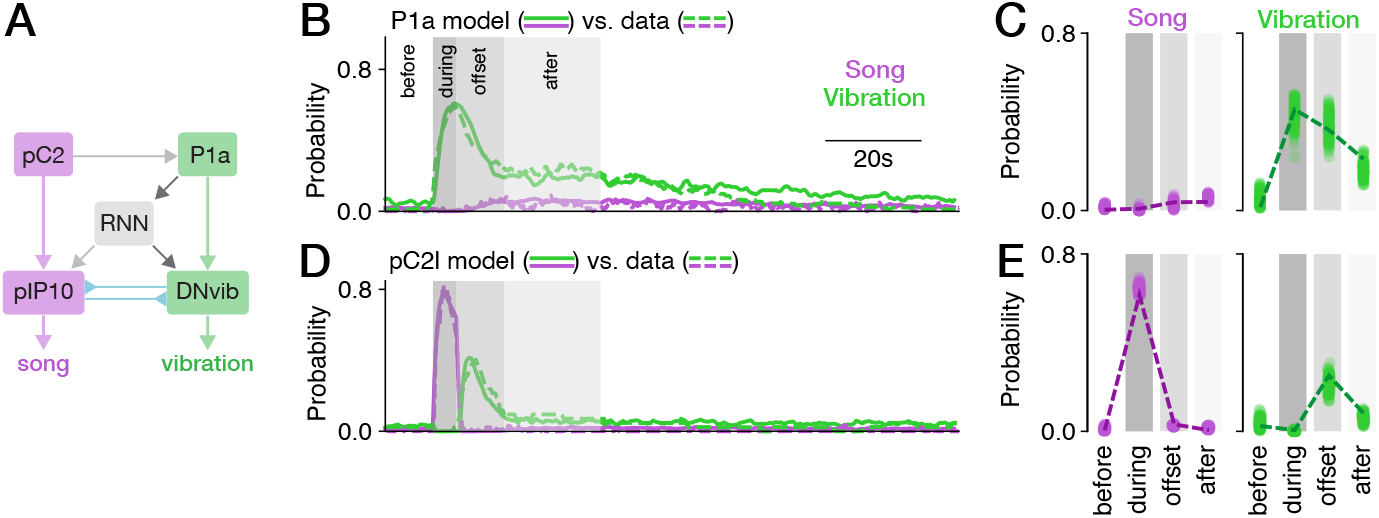
A neural circuit model proposes elementary computations underlying multimodal signaling. **A** Network diagram of the circuit model. Regular and inverted arrows heads indicate excitatoryand inhibitory connections, respectively. **B, D** Song (purple) and vibration (green) for activation of P1a (B) and pC2l (D) in the model (solid lines) and the data (dashed lines, data from Figs 4C, E). The model reproduces the data well: The mean-squared error between model and data is <0.003 for all traces. **C, E** Probability of observing song (purple) and vibration (green) in different epochs around the activation of P1a (C) and pC2l (E) in the model (dots correspond to model runs with independent noise) and the data (dashed lines, data from Figs 4D, F).

This model successfully reproduced the behavioral dynamics. Activating the model P1a neurons produced vibration, followed by a persistent phase of mainly vibration and only little song, that both decay over time (Fig. 6B, C). Activation of pC2l neurons in the model yielded song, directly followed by vibrations (Fig. 6D, E, S9A–C). The persistent phase was mediated by the RNN (Fig. S7). Ablation of the RNN nearly abolished signals after P1a neuron activation during the persistent phase, but did not strongly affect the offset vibrations evoked by pC2l neuron activation and mediated via the slowly decaying dynamics intrinsic to P1a neurons (Fig. S9D–F). Mutual inhibition was required in the model to reduce the overlap between song and vibration, as in natural courtship (Fig. S2, 5C–F), and in the model, vibrations were suppressed when pC2l neurons were activated and song was suppressed when P1a neurons were activated (Fig. S9G–I). The circuit model also reproduced motivational effects in the circuit (Fig. 5I–N). Reducing the excitability of pC2l neurons, P1a neurons, and the recurrent network, reduced song during pC2l neuron activation and strongly reduced the vibrations after activation of pC2l or P1a neurons (Fig. S9J–L). This neural circuit model replicated our behavioral findings and therefore provides insights into the circuit mechanisms that coordinate multimodal signaling behaviors.

## Discussion

We have identified the behavioral contexts (Fig. 2, 3) and circuit motifs that drive multimodal communication signals in *Drosophila* males (Fig. 4, 5, 6). This circuit generates signals with long-lasting, cell-type specific dynamics (Fig. 4, 5), sets the locomotor state required for efficient signal transmission (Fig. 2G–I, 4G–K), and controls both signals through motivational state (Fig. 5H–N).

We found that males produce vibrations when stationary (Fig. 2, 3), a context that previous studies interpreted as an idle state [9, 63]. We show that males are not necessarily idle when sitting next to the female but actively produce communication signals, highlighting the importance of recording all behaviors for correctly interpreting behavioral contexts and the underlying neural circuits [39]. By vibrating primarily when he and the female are stationary and thus when the sender’s and receiver’s legs have full contact with the substrate, the male improves the transmission of vibrations: Vibrations are transmitted via the legs to the substrate, since the abdomen moves but does not touch/tap the substrate [5, 73], and they are detected by leg mechanosensors in the female [41]. Walking therefore interferes with the transmission and detection of vibrations. Song on the other hand is airborne and it’s transmission is not impaired by walking (Fig. 2G–I). But since the song is detected using a highly directional sound receiver [74], it is produced at a more restricted set of positions (Fig. S3E, F). The P1a neurons drive vibrations and induce male stationarity and therewith a locomotor state that favors the transmission of the vibrations (Fig. 4B–D, 4G–K). This coordination of signaling with ongoing behaviors like locomotion or respiration to optimize signal transmission is a general principle of behavioral control. For instance, vocalizations and respiration are coordinated in birds or mammals through shared circuits [34, 35, 75].

Female stationarity was previously [5, 41, 76] interpreted as the effect of vibrations while our behavioral analyses (Fig. 2) and interventions (Fig. 3) show that it is the cause: Stopping the female during courtship is sufficient to drive male vibrations. Both findings can be reconciled: Song, often produced when the male chases the female, slows and stops her [30, 77, 78]. Vibrations, being produced when the female is stationary (Fig. 2) [5], might then prolong phases of stationarity. More experiments will be necessary to elucidate the behavioral effects of song and vibration and to identify the circuits that process both signals [76, 79, 80].

Multimodal signals are driven by an integrated neural circuit in *Drosophila*: The P1a and pC2l neurons—previously considered “song neurons”—drive song and vibration with complex and persistent dynamics (Fig. 4). Multimodal signaling via a single circuit is likely a general principle, since it facilitates signal coordination and modulation (Fig. 5). The periaqueductal gray (PAG) is hypothesized to control multimodal signaling in mammals and birds and shares properties with the proposed circuit in *Drosophila* [1]: The PAG drives vocalizations [29, 81], integrates contextual and motivational information, and innervates multiple premotor regions that control different motor programs [1]. However, precise circuit interactions that might control multimodal signaling in the PAG remain to be identified.

We propose elemental motifs that coordinate multimodal signaling in *Drosophila* using genetic manipulations combined with a computational model. First, direct connections between P1a and pC2l and descending neurons allow external sensory cues to directly and rapidly affect signaling (Fig. 2, 5A–F, S6). Visual motion cues from the walking female activate pC2l [10, 60] to drive song when the male and/or the female move. Notably, song slows the female [30, 78], thereby creating the behavioral context for vibrations. The song-vibration sequence evoked by optogenetic activation of pC2l (Fig. 4E) may therefore constitute a motor prior that facilitates this signal sequence. P1a activity is controlled via chemosensory inputs [46] but the specific cues that drive vibrations in P1a are unclear. The male is too far from the female for contact pheromones (Fig. S3E, F) but volatile pheromones re-activating P1a neurons in an aroused male might suffice [82].

Our experiments also showed that slow dynamics and recurrence act as a memory of the female cues and enable persistent courtship signaling in the absence of constant input from interaction partners (Fig. 4C–F, [83]). These motifs are also found in other systems and therefore likely constitute universal building blocks for controlling behavior: For instance, recurrent circuits in the ventromedial nucleus of the hypothalamus (VMHvl) of mice are central to generating persistent social behaviors that can be easily manipulated by sensory cues through line attractor dynamics [37, 84, 85]. While elucidating the precise circuit, cellular, and molecular mechanisms underlying these common dynamics is challenging in vertebrates models, it will be much more feasible in *Drosophila* given that we have genetic access to identified cell types and connectomics [71].

Lastly, mutual inhibition downstream of P1a and pC2l— between the DNs (Fig. 5A–F, 6) or downstream in the VNC—coordinates multimodal signaling at the motor level to prevent the overlap between song and vibration (Fig. 1H). Mutual inhibition is a core motif whenever mutually exclusive behaviors or patterns of muscle activity are produced by the nervous system—during perceptual decision making, action selection, or motor pattern generation [67, 68, 86, 87].

The descending pathways by which P1a controls locomotor state and vibrations remain to be identified. Unlike pulse and sine, which occur in complex bouts with rapid mode switches [10], direct/immediate transitions between song and vibration are rare during courtship (Fig. 1I, S2). Accordingly neither pMP2 nor pIP10 drive vibrations (Fig. 4B) and vibrations are likely driven by an unknown DNvib (Fig. 6). The complete wiring diagrams of the male brain and VNC will facilitate the identification of descending pathways and pattern generating circuits downstream of P1a that control multimodal signaling and locomotor state in *Drosophila* [71, 88–90]. Ultimately, vibrations are likely produced by thoracic and abdominal contractions that are transmitted via the legs to the substrate [91]. The thoracic muscles, which include the wing muscles that are also required for singing [92, 93], may therefore also contribute to vibrations [73] and may constitute, after the divergence of pathways at the premotor level, a convergent *final common pathway* [94] for multimodal signaling in *Drosophila*.

Overall, our results identify common circuit motifs—feedforward excitation, recurrence, mutual inhibition—that can be combined in a single circuit to support dynamical and context-specific multimodal signaling. Moreover, we establish *Drosophila* as a new model system for studying multimodal communication.

## Acknowledgments

We thank Tina Zahrie, Jannis Hainke, Maximilian Ferle, Karla Rivera, Alina Seidel for help with annotation and data acquisition, Frank Kötting, Stephan Löwe from the ENI workshop for help with designing behavioral chambers, Gesa Hoffmann, Jan Schöning, Christine Gündner, Christiane Becker for technical and adminstrative assistance. Martin Göpfert und Philip Hehlert provided access to a laser vibrometry setup. Gwyneth Card, David Anderson, Vivek Jayaraman, André Fiala, Peter Andolfatto, Joshua Lillvis, Martin Göpfert, Janelia flylight, Bloomington stock center for gifts of flies. We thank all members of the Clemens lab as well as Frederic Roemschied, Daniela Vallentin, Mala Murthy, and Xinping Li for feedback on the manuscript. This work was funded via an Emmy Noether Grant (Project number 329518246) and an ERC Starting Grant (Grant agreement No. 851210) to JC.

## Author contributions

- Conceptualization - ES, AK, JC
- Animals and behavioral experiments - ES, AK, MS, BS SR, KA
- Modeling and analysis - ES, JC
- First draft - ES, JC
- Feedback on draft - AK, MS, BS SR, KA

## Methods

### Fly strains and rearing

Flies were kept on a 12:12 hour dark:light cycle, at 25°C and 60% humidity. Flies were collected as virgins within 8 hours after eclosion, separated by sex, and then housed in groups of 3-15 flies.

### Behavioral setups

The behavioral chamber measured 44 mm in diameter and 1.9 mm in height; chamber and lid were machined from transparent acrylic. Chamber lids were coated with Sigmacote (Sigma-Aldrich) to prevent flies from walking on the ceiling, and kept under a fume hood to dry for at least 10 minutes.

The floor of the chamber was tiled with 16 microphones (Knowles NR-23158) that were embedded into a custom-made PCB board (design modified from Coen et al. [8]). The microphones were covered with a thin, white paper for the flies to walk on and to record sound and vibration. Microphone signals were amplified using a custom-build amplifier [49] and digitized using a data acquisition card (National Instruments Pcie-6343) at a sampling rate of 10 kHz.

Fly behavior was recorded from above using a USB camera (FLIR flea3 FL3-U3-13Y3M-C, 100 frames per second (fps), 912 × 920 pixels), equipped with a 35 mm f1.4 objective (Thorlabs MVL35M1). The chamber was illuminated with weak blue light (470 nm) and white room light. For optogenetic experiments, the room light was turned off, to reduce interference between illumination and activation wavelengths. A 500 nm shortpass filter (Edmund Optics, 500 nm 50 mm diameter, OD 4.0 Shortpass Filter) filtered out green (525 nm) and red (625 nm) wavelengths used for optogenetics.

To match the males’ abdominal quivering with the vibration pulses recorded on the microphones, we recorded videos with higher spatial (1200 × 1200 pixel frames covering a chamber with diameter 11 mm) and temporal (150 fps) resolution. The chamber was centered on one of the 16 recording microphones and illuminated with white LEDs.

Synchronized recordings of audio, video, and delivery of optogenetic stimuli was controlled using custom software https://janclemenslab.org/etho.

As a control, we also measured the substrate deflections induced by vibrations using a PSV-400 laser Doppler vibrometer (Polytec GmbH) in the same chamber and paper substrate used above. The laser beam was directed through the transparent lid perpendicular to the paper surface at a distance of 1-4 mm near a stationary male courting a female (Fig. S1). Data obtained with the laser vibrometer were high-pass filtered (Butterworth, 60 Hz) before analysis.

### Behavioral assays

For all experiments, 3 to 7 day old naive males and virgin females were used. Flies were introduced gently into the chamber using an aspirator. All recordings were performed during the flies’ morning activity peak and started within 120 minutes of the incubator lights switching on. Recordings of video and audio were performed for 30 minutes in the regular chamber, for 10 minutes in the smaller chamber, and for 2 minutes during laser vibrometry.

In experiments using males with amputated wings (Fig. S1G–H), flies were cold-anesthetized and both wings were cut using fine scissors at least 18 hours before the experiment.

To induce sexual satiation (Fig. 5H–N) males were transferred individually into food containing vials with 10-15 virgin NM91 females and allowed to freely interact and copulate for 4-6 hours. The control males came from groups of 10-15 males with the same genotype (pC2l-CsChrimson or P1a-CsChrimson). After the pre-exposure period, all flies were quickly anesthetized on ice to separate one male from the group, who was gently transferred into an empty vial to recover for 15 minutes. Then he was gently introduced into the behavioral chamber and the optogenetic activation experiment was started.

### Optogenetics

Flies were kept for at least 3 days prior to the experiment on food containing retinal (1 ml all-trans retinal (Sigma-Aldrich) solution (100 mM in 95% ethanol) per 100 ml food). To prevent the degradation of the retinal and continuous neural activation, the vials were wrapped in aluminium foil. Control flies were either parental controls (Fig. 3, 4) or had the same genotype as experimental flies and were kept on regular food without additional retinal. Note that regular food contains trace amounts of retinal, and drivers with strong expression can therefore produce effects even in the non-retinal controls.

For neural inactivation, we used the GtACR1 channel [53, 96], which was excited using a green LED (625 nm). For inactivation of vGlut (Fig. 3A–B) we used an LED intensity of 14 mW/cm^2^. Experiment consisted of 40 trials of optogenetic stimulation. Each trial started with 5 s stimulation (green LED on) followed a pause of 25 s. For neural activation, we used the CsChrimson channel [95], which was activated using a red LED (625 nm). For activation of DNp28 (Fig. 3C–D) we used an LED intensity of 89 mW/cm^2^. Each experiment consisted of 30 trials of optogenetic stimulation. Each experimental trial started with 5 s stimulation followed by a pause of of 25 s. For pC2l and P1a activation (Fig. 4–5) we used LED intensities 14, 27, 83, and 209 (P1a only) mW/mc^2^. Each experiment consisted of 7 trials of optogenetic stimulation and each trial started with 5 s of optogenetic stimulation followed by pause of 120 s.

### Analysis of microphone signals

Multimodal courtship signals (pulse, sine, vibration) were manually annotated using the graphical user interface of DAS [97]. For optogenetic manipulation of female walking (Fig. 3) and the satiation assay (Fig. 5H–N), the annotators were blind to experimental condition.

*Pulse and vibration trains* were defined as groups of pulses with an interval less than 2–2.5 the modal interval (80 ms for pulse song, 400 ms for vibration). The *signal fraction* is the fraction of all courtship frames in which a specific signal—pulse, sine, or vibration—was produced.

*Transition probabilities* between signals correspond to the fraction of signals of a given type that were followed by a given other signal (i.e. fraction of pulse trains followed by sine song, or pulse song, or vibrations), regardless of the duration of the silent pause between trains. We then averaged the transition probabilities over all 14 pairs of NM91 wild type flies.

*Signal probabilities* for experiments with optogenetic neural activation or inactivation, are given as the fraction of trials during which sine song or pulse and vibration trains were produced. We then computed the mean across trials pooled across all males. For experiments with speed-controlled females (Fig. 3) and with optogenetic activation of P1a and pC2l in males paired with a female (Fig. 5C–F), we only considered time points during which the male courted the female.

### Behavioral data analysis

Flies were tracked using standard procedures (estimation of background as median frame, subtraction of background from each frame, thresholding, localization of flies using Gaussian mixture model). The location of individual body parts (head, thorax, abdomen, left and right wing) were then tracked using DeepPoseKit [98]. For most analyses, the tracking data was downsampled from the original frame rate of 100 Hz (fps) to 50 Hz. All time points after the beginning of copulation were excluded from analysis.

To show traces of signal probabilities or velocities for optogenetic experiments or onset/offset analysis (Fig. 3–5, S4–5), we pooled data across flies and computed the mean (for signal probabilities) or median (for velocities) across stimulation trials or onsets and offsets. To eliminate tracking errors from velocity or wing angle data, we excluded data points where the distance between male and female thoraces dropped below 1 mm and were the tracking confidence for the head or thorax was less than 50%. All traces shown for optogenetic experiments (Fig. 3–5) are smoothed with a Gaussian window with a standard deviation of 0.1 s.

*Courtship* was defined as time points during which the male was within 8 mm (6 mm for GLM analysis) of the female and ±60° behind her. The *courtship index* is the fraction of time points that are courtship from the beginning of the recording until copulation started or the recording ended.

### Correlating abdominal quivering and vibration pulses

Flies positions and body parts in the high-resolution videos (150 fps, 1200 × 1200 pixels at 11 mm) were tracked using SLEAP [50]. We then independently annotated abdominal quivering in the video, visible in the top-down view as a brief shortening of the abdomen, and vibration pulses in the audio.

### Behavioral modeling

Multinomial Generalized Linear Models (GLMs) were used to identify the behavioral cues and contexts that drive the choice between song (pulse, sine) and vibration. Models were fitted to predict whether the male produced song, vibration or no signal at any moment in time.

As behavioral cues, we extracted 19 metrics from the fly tracks of 14 male-female pairs of NM91 using xarray-behave (Table 5): male or female rotational speed, rotational acceleration, velocity and its forward and lateral components, acceleration and its forward and lateral components, male-female distance, as well as the male’s relative angle (male position relative to female body axis) and relative orientation (males heading relative to female center). We only considered courtship frames and frames before copulation.

**Table 1:**
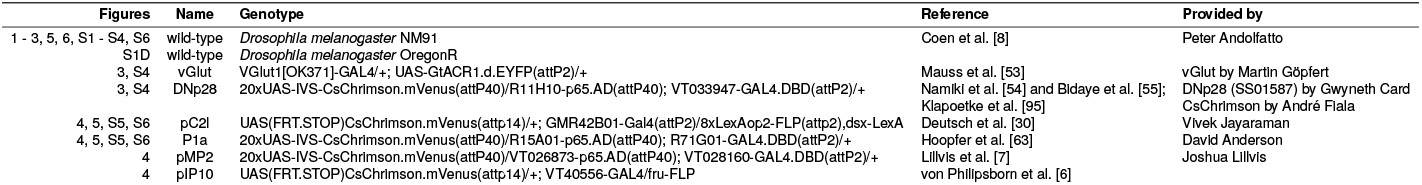
Fly lines used.

**Table 2:**
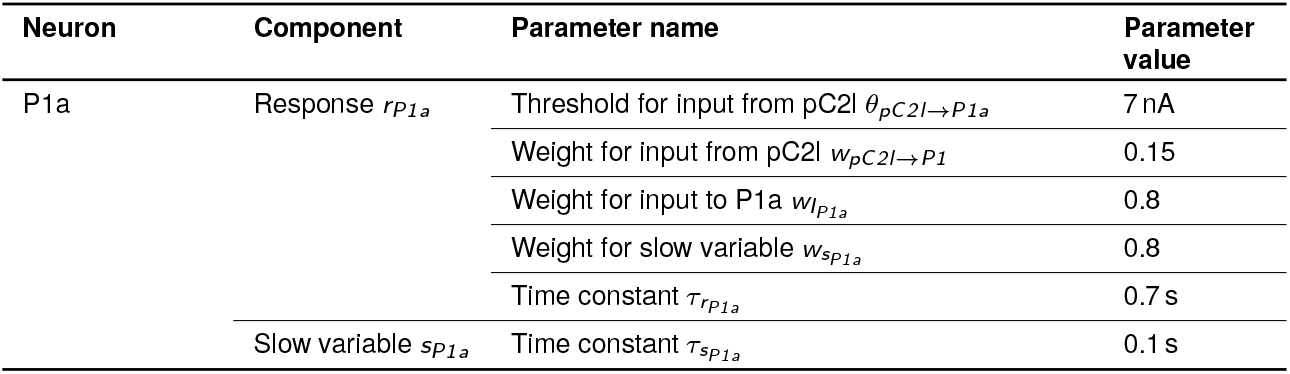
Model parameters for P1a.

| Neuron | Component | Parameter name | Parameter value |
| --- | --- | --- | --- |
| P1a | Response $r_{P1a}$ | Threshold for input from pC2I $\theta_{pC2I \rightarrow P1a}$ | 7 nA |
| | | Weight for input from pC2I $w_{pC2I \rightarrow P1}$ | 0.15 |
| | | Weight for input to P1a $w_{I_{P1a}}$ | 0.8 |
| | | Weight for slow variable $w_{s_{P1a}}$ | 0.8 |
| | | Time constant $\tau_{r_{P1a}}$ | 0.7 s |
| | Slow variable $s_{P1a}$ | Time constant $\tau_{s_{P1a}}$ | 0.1 s |

**Table 3:** Model parameters for the recurrent neural network (RNN).

| Neuron | Component | Parameter name | Parameter value |
| --- | --- | --- | --- |
| RNN | Inputs $I_{RNN}$ | Threshold for input from P1a $\theta_{P1a \rightarrow RNN}$ | 1.6 nA |
| | | Time constant $\tau_{I_{RNN}}$ | 16 s |
| | Response $r_{RNN}$ | Weight for recurrence $w_{p_{RNN} \rightarrow r_{RNN}}$ | 0.96 |
| | | Time constant $\tau_{r_{RNN}}$ | 0.7 s |
| | Recurrence $p_{RNN}$ | Time constant $\tau_{p_{RNN}}$ | 2 s |

**Table 4:** Model parameters for pIP10 and DNvib.

| Neuron | Component | Parameter name | Parameter value |
| --- | --- | --- | --- |
| pIP10 | Response $r_{pIP10}$ | Weight for input from RNN $w_{RNN \rightarrow pIP10}$ | 1.6 |
| | | Weight for mutual inhibition from DNvib $w_{m_{DNvib}}$ | 10 |
| | Nonlinearity $\Sigma_{pIP10}$ | Saturation $\omega_{pIP10}$ | 20 |
| DNvib | Response $r_{DNvib}$ | Weight for input from P1a $w_{P1a \rightarrow DNvib}$ | 1.5 |
| | | Weight for input from RNN $w_{RNN \rightarrow DNvib}$ | 1.92 |
| | | Weight for mutual inhibition from pIP10 $w_{m_{pIP10}}$ | 1.5 |
| | Nonlinearity $\Sigma_{DNvib}$ | Saturation $\omega_{DNvib}$ | 3 |
| pIP10 or DNvib | Response $r_{pIP10}$ or $r_{DNvib}$ | Time constant $\tau_r$ | 1 s |
| | Adaptation $a_{pIP10}$ or $a_{DNvib}$ | Time constant $\tau_a$ | 5 s |
| Mutual inhibition from DNvib or pIP10 | Response $m_{pIP10}$ or $m_{DNvib}$ | Weight for input from DNvib or pIP10 $w_r$ | 0.001 |
| | | Time constant $\tau_m$ | 1 s |
| | Adaptation $a_{m_{pIP10}}$ | Time constant of adaptation $\tau_{a_{m_{pIP10}}}$ | 1 s |
| | | Weight for input from adaptation $w_{a_{m_{pIP10}}}$ | 10000 |

**Table 5:** Open source software used.

| Resource | Link (citation) |
| --- | --- |
| DeepPoseKit | https://github.com/jgraving/DeepPoseKit [98] |
| DeepAudioSegmenter | https://github.com/janclemenslab/das [97] |
| GLM utilities | https://github.com/janclemenslab/glm_utils |
| Inkscape 0.92 | https://inkscape.org |
| Python 3.7–3.12 | https://python.org |
| scikit learn | https://scikit-learn.org [100] |
| seaborn | https://seaborn.pydata.org [111] |
| SLEAP | https://sleap.ai [50] |
| xarray-behave | https://github.com/janclemenslab/xarray-behave |
| etho | https://github.com/janclemenslab/etho |
| pandas | https://pandas.pydata.org [112] |
| numba | https://github.com/numba/numba [113] |
| networkx | https://networkx.org [108] |
| navis | https://navis-org.github.io/navis [109] |
| natverse flybrains | https://natverse.org [114] |
| flywire codex | https://codex.flywire.ai [104] |

The cues for each pair were z-scored and then pooled across pairs. That way, each GLM was fitted to the data from multiple pairs. Since we were interested in identifying the time course of each cue that best predicted signaling, we delay-embedded the cues. That is, the signals in each time point was predicted using the time course of each cue in the 1 s preceding that time point. To reduce dimensionality, we projected each 1 s onto a basis of four raised cosines covering the 1 s time window with logarithmic spacing [99]. Thereby, the cues’ time course in the 1 s preceding each time point was predicted by 4 values. The temporal filters (Fig. 2H) were recovered from the 4 weights learned by the GLM by back-projecting the raised cosine basis to time. The filter sum (Fig. 2G, S3B), was given by the sum of all filter values in the time domain.

Since the fraction of song, vibration and no signal in the data were skewed towards no signals, we balanced the data prior to fitting, by randomly sub-sampling an equal number from each prediction target (song, vibration, no signal). This yielded 73,562 time points per signal type as inputs to the models fitting.

#### GLM fitting and evaluation

Data points of behavioral cues were split into 90% training data and 10% test data. Each model was fitted 10 times, each time with randomly train-test splits and balancing. Models were fitted using LogisticRegressionCV from scikit-learn [100], with L2 regularization, ten-fold cross-validation and a maximum of 500 iterations.

The performance of each fitted model was quantified by comparing model predictions on the test set to behavioral groundtruth data. Predicted and true signals were tabulated in a confusion matrix, normalized by the true signals (Fig. 2C, E). Diagonal matrix elements correspond to correct predictions (plotted in Fig. 2F) and off-diagonal elements correspond to prediction errors. To obtain a single score of the performance, we computed the accuracy as the average over the diagonal values. We fitted two types of models to assess the contribution of individual cues to the males’ signal choice. To assess the general ability of the cues to predict the males’ signal choice, we fitted a model that used all 19 cues (Fig. 2C). As a second step, to assess to information contributed by each individual cue, we fitted a separate models for each cue and assessed their performance (Fig. 2D).

#### Connectome analyses

Connectome analyses in Fig. S8 were based on the female whole brain connectome, flywire [101–103], since no male brain connectome data is currently available. The data was downloaded from flywire codex (https://codex.flywire.ai/api/download, v783) [104] and further processed using open source packages (see Table 5). pC1 and pC2 neurons were identified based on existing cell-type annotations in flywire [103] and connections [105–107] were identified using the all_simple_paths function of the networkx package [108]. The outline of the brain and the neuronal skeletons were plotted using navis [109] and natverse’s flybrains package [110].

### Circuit model

#### Model structure and working principle

The primary goal of the model is to synthesize the experimental results and show that our current model of the circuit is sufficient to explain the behavioral data. The model is well supported by existing and our own data, and consists of four main components:

1. The social cue integrating neuron groups P1a and pC2l mediate acute effects of activation via connections to descending command-like neurons.
2. A recurrent neural network (RNN) downstream of P1a mediates the long-term effects of circuit activation.
3. Two descending command-like neurons, pIP10 and DNvib, drive song and vibration in the ventral nerve chord.
4. Mutual inhibition between or downstream of pIP10 and DNvib reduces the overlap between song and vibration.

P1a and pC2l have been shown to be activated by social cues in numerous studies. The pC2l neurons are activated by male pulse song [30] and likely also visual [60] and other cues. The P1a neurons receive inputs from volatile and contact chemical cues [46, 61, 62]. Our behavioral results leave open the possibility that additional, still unidentified cues activate P1a.

In our experiments, activation of P1a and pC2l drove vibration and song, respectively, with short latency (Fig. 4). This suggest that they have short connections spanning only one or a few synapses to command-like descending neurons. Direct connectivity between pC2l and the song DN pIP10 has been established anatomically and functionally [59]. Short connections between P1a and descending command neurons are not known but are likely given the behavioral data. This connection can be tested directly once DNvib has been identified.

Vibrations were also driven at the offset of pC2l. In the model, this is mediated via a pC2l to P1a connection (Fig. S7B, E). pC2l activity would induce relatively weak and slowly decaying activity in P1a. A pC2l to P1a connection has been hypothesized in a recent paper on song patterning [10] and was required to explain the production of complex song upon pC2l activation. Our data provides independent support for such a connection. The activity of P1a has been shown to decay slowly with a time constant of 5–10 s [62] which matches the time constant of the offset vibrations after pC2l activation (Fig. 4). This supports the idea of offset vibrations after pC2l activation being driven by this slowly decaying P1a activity.

An RNN downstream of P1a maintains vibration activity for tens of seconds. Elements of the RNN have been characterized previously using behavioral and imaging experiments, and the pCd neurons are members of this network [36]. Connectivity downstream of the RNN is unknown. For simplicity, we assume that the RNN drives both song and vibration DNs. However alternative implementations are possible. Signaling after P1a activation in solitary males is strongly biased towards vibrations and this is reflected in stronger relative connectivity from the RNN to the DNvib versus pIP10 in our model.

Lastly, mutual inhibition downstream of P1a and pC2l reduces the overlap between song and vibration, and induces switching between song and vibration during the persistent phase driven by adaptation and noise. This component of the model is derived from models of bistable phenomena [68]. Mutual inhibition could be implement at different stages downstream of P1a and pC2l: Upstream of pIP10 and DNvib, between pIP10 and DNvib, or downstream of the DNs in the VNC. For simplicity, we model mutual inhibition as happening between pIP10 and DNvib. pIP10 receives input from pC2l and the RNN, and DNvib receives input from P1a and the RNN. Both DNs adapt, which is supported by the observation of spike-frequency adaptation in patch clamp recordings of pIP10 [10]. pIP10 activity drives song in the VNC and an interneuron that inhibits DNvib. DNvib activity drives vibrations in the VNC and an interneuron that inhibits pIP10. The latter interneuron adapts, which acts as a high-pass filter that speeds up the inputs from P1a-DNvib to account for the short latency of inhibition of song upon P1a activation (Fig. 5). Gaussian noise is added to the output of pIP10 and DNvib to enable stochastic switching between song and vibration in the persistent phase.

Since we were interested in circuit dynamics on a timescale of seconds, we implemented the a rate-based model, in which the activity of individual neurons is represented by continuous variables that are considered to be proportional to the firing rate of the cell (individual cells, e.g. for pIP10, or cell clusters, e.g. P1a or pC2l). To translate the activity of pIP10 and DNvib to behavior, we consider their activity to be proportional to the probability of observing song and vibration, respectively. Trial averaged plots show the average probability over 100 model simulations with different noise patterns.

#### Mathematical details

##### pC2l

The population activity of the pC2l neurons is a copy of their optogenetic input: *r*_*pC2l*_ = *I*_*opto* → *pC2l*_. Optogenetic input was modeled as rectangular pulses with the same duration as used in the experiments (5 s, interleaved by a pause of 120 s). We assumed a logarithmic mapping from LED intensity to input current (14, 27, 42, 83 mW/cm^2^ -> 0.5, 0.6, 1.1, 1.4 nA).

##### P1a

The inputs to P1a are given by:

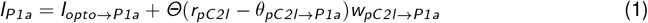

where *I*_*opto*→ *P1a*_ is the input from optogenetic activation (or sensory cues), and *r*_*pC2l*_ is the input from pC2l which is passed through a threshold-linear function

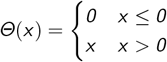

after subtraction of a threshold term *θ*_*pC2l*→ *P1a*_. The threshold ensures that weak activation of pC2l is insufficient to drive offset vibrations via P1a (Fig. S5F). As for pC2l, we assumed a logarithmic mapping from LED intensity to input current (14, 27, 42, 83 mW/cm^2^ -> 0.12, 0.16, 0.20, 0.24 nA). The response of P1a is given by

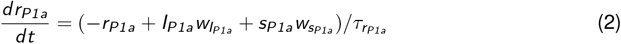

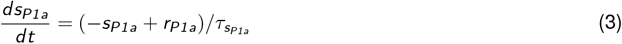

where *r*_*P1a*_ is a continuous variable proportional to the population firing rate of the P1a neurons, *I*_*P1a*_ are the external inputs to P1a (Eq. 1) with weight 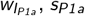 is the input from a slow variable with weight 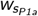, and 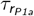 is the membrane time constant. The slow decay of P1a activity [62] is replicated by a positive feedback loop between *r*_*P1a*_ and a slow variable, *s*_*P1a*_. The slow variable could represent cell-intrinsic mechanisms arising from slow calcium dynamics coupled with calcium-activate sodium channels. The slow variable receives input from P1a, *r*_*P1a*_, and is integrated with time constant 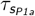. Before being passed on to downstream partners, the output of P1a is transformed using a static logarithmic nonlinearity to mimic response saturation *r*_*P1a*_ = log(*1* + *2 r*_*P1a*_).

#### Recurrent neural network

While the slow variable, *s*_*P1a*_ (Eq. 3), reproduces the known slow decay of P1a activity [62], a recurrent neural network (RNN) downstream of P1a generates persistent signaling over tens of seconds after P1a activation [36]:

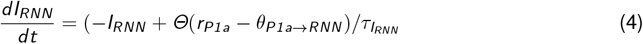

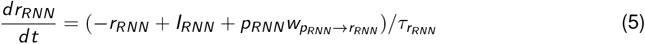

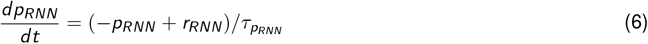

External input to the RNN, *I*_*RNN*_, from P1a is passed through a threshold-linear function with threshold *θ*_*P1a*→ *RNN*_ and integrated with time constant 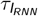. The threshold ensures that only strong activation of P1a elicits persistence, not the weak activation from pC2l. Input from the recurrent pool, *p*_*RNN*_, is integrated with weight 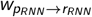 and together with external input, *I*_*RNN*_, integrated with a time constant 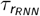. The recurrent pool receives input from the RNN itself and has a time constant 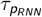.

#### Descending neurons pIP10 and DNvib

The pIP10 neuron integrates input from the RNN and from pC2l, mutual inhibition from DNvib, adaptation, and noise:

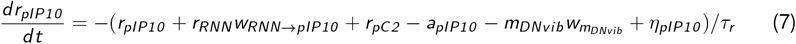

where 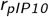 is the activity of pIP10, *r*_*RNN*_ is the input from the RNN with weight *w*_*RNN*→ *pIP10*_, *r*_*pC2*_ is the input from pC2l, *a*_*pIP10*_ is an inhibitory adaptation current (see eq. 9 below), *m*_*DNvib*_ is an inhibitory input from DNvib with weight 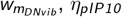 is Gaussian noise (see eq. 10 below), and *τ*_*r*_ is an integration time constant.

Similar to pIP10, DNvib integrates inputs from the RNN and P1a, mutual inhibition from pIP10, adaptation and noise:

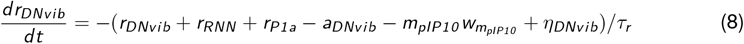

where *r*_*DNvib*_ is the activity of DNvib, *r*_*RNN*_ is the input from the RNN, *r*_*P1a*_ is the input from P1a, *a*_*DNvib*_ is an inhibitory adaptation current, *m*_*pIP10*_ is an inhibitory input from pIP10 with weight 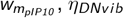 is Gaussian noise (see eq. 10 below), and *τ*_*r*_ is an integration time constant.

To enable bistable dynamics with noise-induced switching between song and vibration after activation of P1a, we added an adaptation current and noise to pIP10 (eq. 7) and DNvib (eq. 8) [68]. The adaptation is modeled as negative feedback:

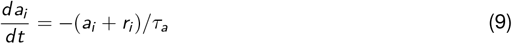

where *a*_*i*_ is the adaption current for neuron *i, r*_*i*_ is activity of neuron *i*, and the adaptation time constant is *τ*_*a*_. Gaussian noise *η* with time constant *τ*_*η*_ and standard deviation *σ*_*η*_ was given by:

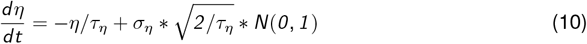

*N*(*0, 1*) is a random variable with zero mean and unit variance.

During integration, *r*_*pIP10*_ and *r*_*DNvib*_ are passed through a nonlinearity *Σ* which limits their activity to an upper bounds of *ω*:

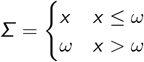

#### Mutual inhibition downstream of pIP10 and DNvib

Mutual inhibition downstream of pIP10 and DNvib is based on a canonical model of bistable perception [68]. In this model, switching arises from adaptation (eq. 9) and noise (eq. 10) in the response of pIP10 and DNvib. We implemented the mutual inhibition via inhibitory interneurons *m*_*DNvib*_ and *m*_*pIP10*_, respectively. Only *m*_*pIP10*_ adapts to speed up the dynamics of the inhibitory inputs from DNvib to pIP10 which are otherwise too slow to mediate strong and fast inhibition of song from DNvib:

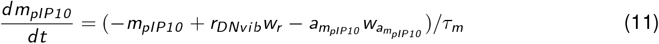

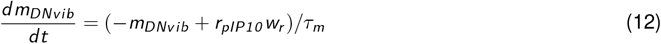

Both *m*_*pIP10*_ and *m*_*DNvib*_ integrate their external inputs with weight *w*_*r*_, and have a time constant *τ*_*m*_. For 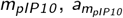 is the adaptation current with weight 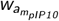 and an adaptation time constant 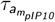 (eq. 9).

#### Model fitting and simulation

The differential equations were solved numerically with the Euler method and a time step of 1 ms, accelerated using just-in-time compilation with numba. The model was fitted by manually adjusting the parameters.

#### Model manipulations

For ablating recurrence (Fig. S9D–F), we set the weights for inputs from the RNN in pIP10 and DNvib, *w*_*RNN*→*pIP10*_ and *w*_*RNN*→*DNvib*_ to zero. For ablating mutual inhibition (Fig. S9G–I) we set the weights for inputs from the mutual inhibition, 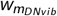 and 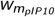 to zero. Effects of sexual satiation in the model (Fig. S9J–L) were reproduced by changing 1) the gain of inputs to pC2l from to 0.6, 2) the weight for the slow variable in P1a, 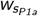, from 0.8 to 0.75, and 3) the weight for recurrent inputs to RNN, 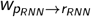, from 0.96 to 0.75.

## Statistical analyses

All tests were Wilcoxon (for paired data) or Mann-Whitney-U test (for unpaired data). The significance levels for multiple comparisons were adjusted from 0.05 using the Bonferroni method. For assessing the effect of optogenetic activation in courting males, statistics only include males that intensely courted the female 10 s before and during optogenetic activation. Intense courtship was defined as a courtship index of 0.9 (see above).

## Supplementary section

**Figure S1:**
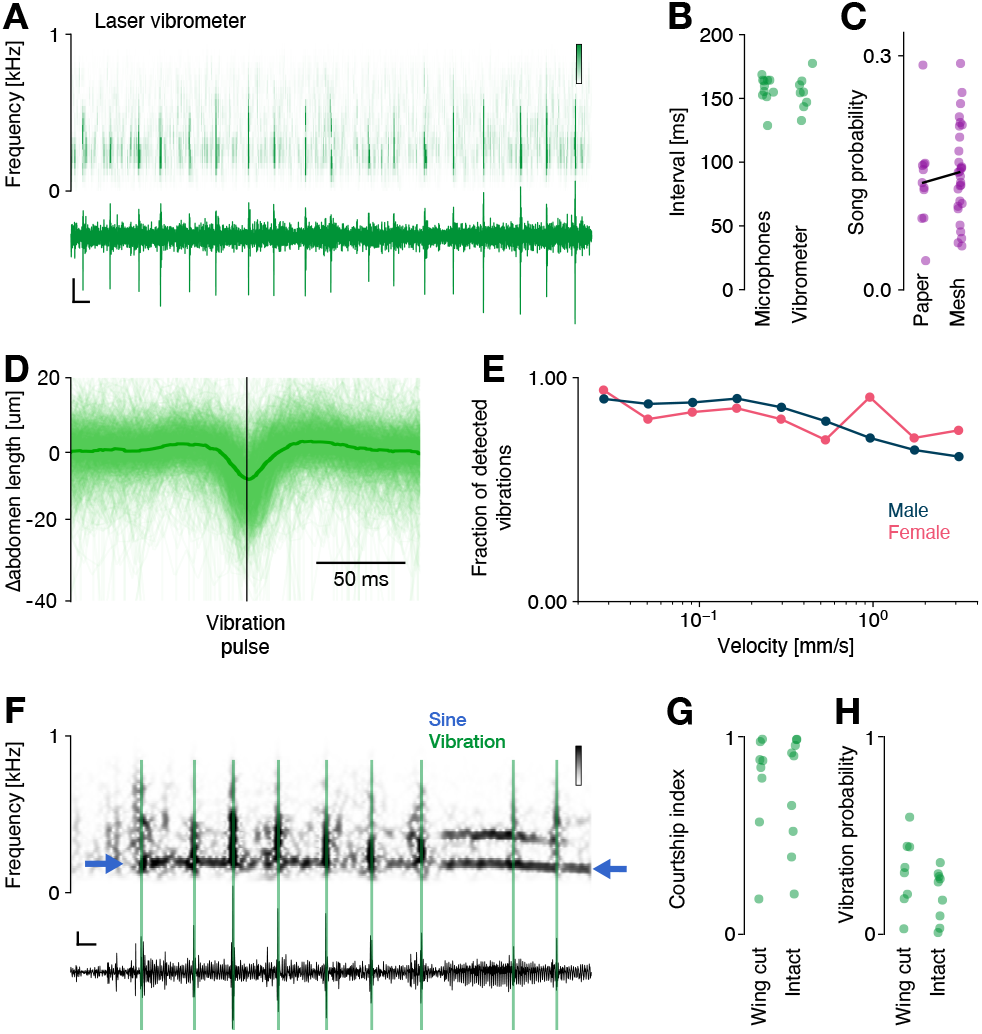
Vibrations can be reliably recorded using a microphone array. **A** Vibrations recorded using a laser vibrometer (bottom) and the corresponding spectrogram (top). Vertical and horizontal scale bar corresponds to 20 nm/s and 100 ms. **B** Intervals between vibrations recorded using laser vibrometry (155±13 ms, N=8 flies) and microphones (160±11 ms, N=11 flies) are similar (p=0.40, two-sided Mann-Whitney U test). Dots correspond to the median vibration intervals of individual males. Intervals between vibration trains (>360 ms) were excluded. **C** Probability of song during courtship recorded in the same 16-microphone chamber with paper (13.7±0.5% (median±IQR), N=11 pairs) and mesh (15.1±0.9%, N=29 pairs) substrates (p=0.61, two-sided Mann-Whitney test). **D** Length of the abdomen extracted from SLEAP tracked male poses aligned to vibration pulses detected on the microphones. Individual vibration pulses are associated with abdominal quivering [5], resulting in a transient shortening of the abdomen. The abdomen length was calculated as the distance between the thorax center and the tip of the abdomen. Individual green lines show individual vibrations, the thick green line is the average over N=747 vibrations. **E** Probability of detecting vibration within 0.1 seconds of male quivering as a function of male (blue) and female (pink) velocity. We binned velocities into 9 logarithmically spaced bins between 0.2 and 2 mm/s and calculated the fraction of detected vibrations. Over all bins, detection probability is at or above 0.80. Thus, the recording system enables reliable recoding of vibrations in stationary and walking flies. **F** Microphone trace (bottom) and spectrogram (top) showing a rare overlap between sine song (dark vertical bands in the spectrogram) and vibrations (green). Vertical and horizontal scale bar corresponds to 0.1 V and 50 ms. **G** Wing cut males court as much as intact males (courtship index wing cut 0.86±0.25 and intact 0.90±0.27, p=0.78, two-sided t-test). **H** Wing cut males vibrate as much as intact males. Probability of vibration during courtship in wing cut and intact males: 0.32±0.16 and 0.26±0.12 (p=0.14, two-sided t-test, N=8 wing-cut, N=9 intact males).

**Figure S2:**
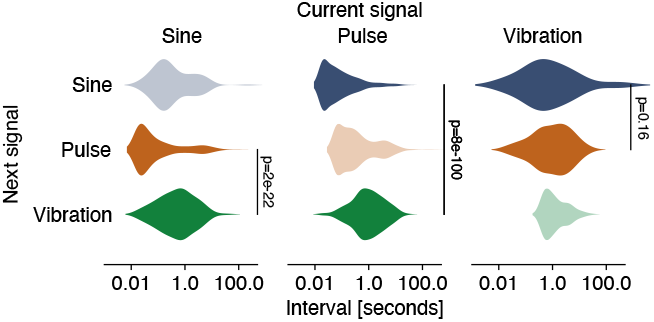
Song and vibrations are temporally separated. Pauses between sine songs, pulse trains, and vibration trains. The song modes are interleaved by much shorter pauses than song and vibration. This is consistent with song and vibration being produced in distinct behavioral contexts (sine to pulse 0.04±0.16 s (median±IQR), pulse to sine 0.06±0.14 s, sine to vibration 0.57±1.16s, pulse to vibration 0.94±1.94 s).

**Figure S3:**
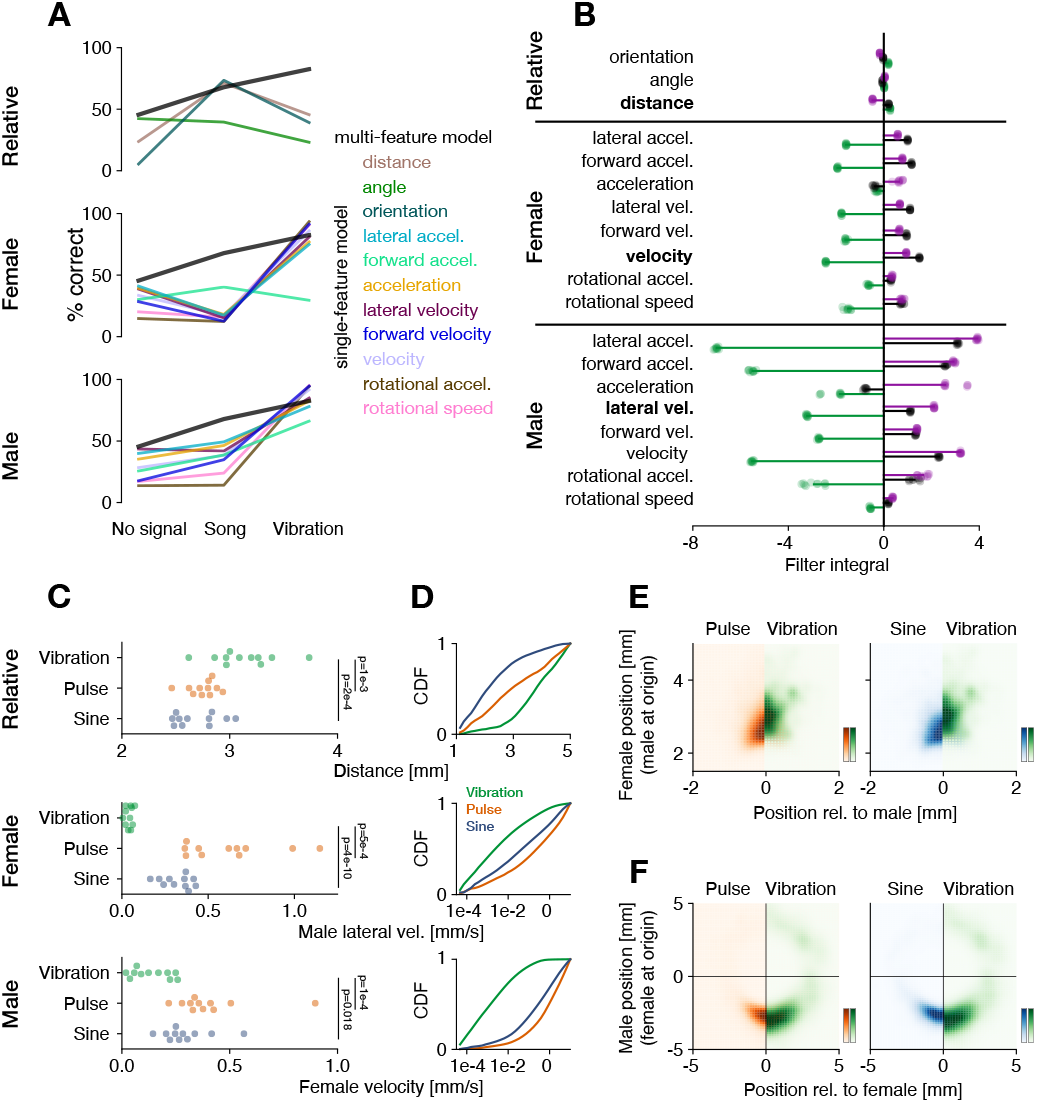
Males vibrate when slow and sing when close to females and when moving. **A** Predictive performance (% correct) of the multi-feature model (black, Fig. 2C) and of the single-feature models (features color coded, see legend) for predicting no signal, song, and vibration. Features are split by their type (relative, female, male). Same data as Fig. 2F, but lines are color-coded by feature. **B** Integral of the linear filters for models fitted with single cues (same models as in Fig. 2D–H). Male and female speed-related cues tend to have filters with negative integrals for vibration (green) and positive integrals for song (purple) and nothing (black). This means that vibrations are mainly produced when flies are slow. Individual dots correspond to the filter integral from 10 fits of the models with independent train-test splits, horizontal lines connect x=0 to the mean over the 10 fits. Same data as Fig. 2G but for all features. **C** Most predictive relative (male-female distance, top), female (velocity, middle), and male (lateral velocity, middle) cues during sine (blue), pulse (orange), and vibration (green). Individual dots show the average value for each of 11 pairs. Distance for sine (2.6±0.3 mm, median±IQR), pulse (2.8±0.1 mm), and vibration (3.1±0.3 mm). The males sing when close to the female and vibrate when further away. Male lateral velocity when producing sine (0.36±0.12 mm/s), pulse (0.62±0.29 mm/s), and vibration (0.05±0.03 mm/s). Female velocity when producing sine (0.27±0.09 mm/s), pulse (0.36±0.09 mm/s), and vibration (0.12±0.15 mm/s). When males or females slow, they tend to vibrate, when they are fast, they tend to sing sine or pulse song. P-values from Dunnet post-hoc tests of a Kruskal-Wallis test (both two-sided). **D** Cumulative density functions of distance (top), male lateral velocity (middle), and female velocity (bottom) for sine (blue), pulse (orange), and vibration (green) (515076 data points of courtship pooled across N=11 pairs). Same data as Fig. 2I but with song split into pulse and sine. **E, F** Position of the female relative to the male (E) and of the male relative to the female (F) for pulse (orange) sine (blue) and vibration (green). Histogram based on the average values positions over whole sine songs or pulse and vibration trains (N=27160/39389/13805 trains or songs for sine/pulse/vibration over N=11 pairs).

**Figure S4:**
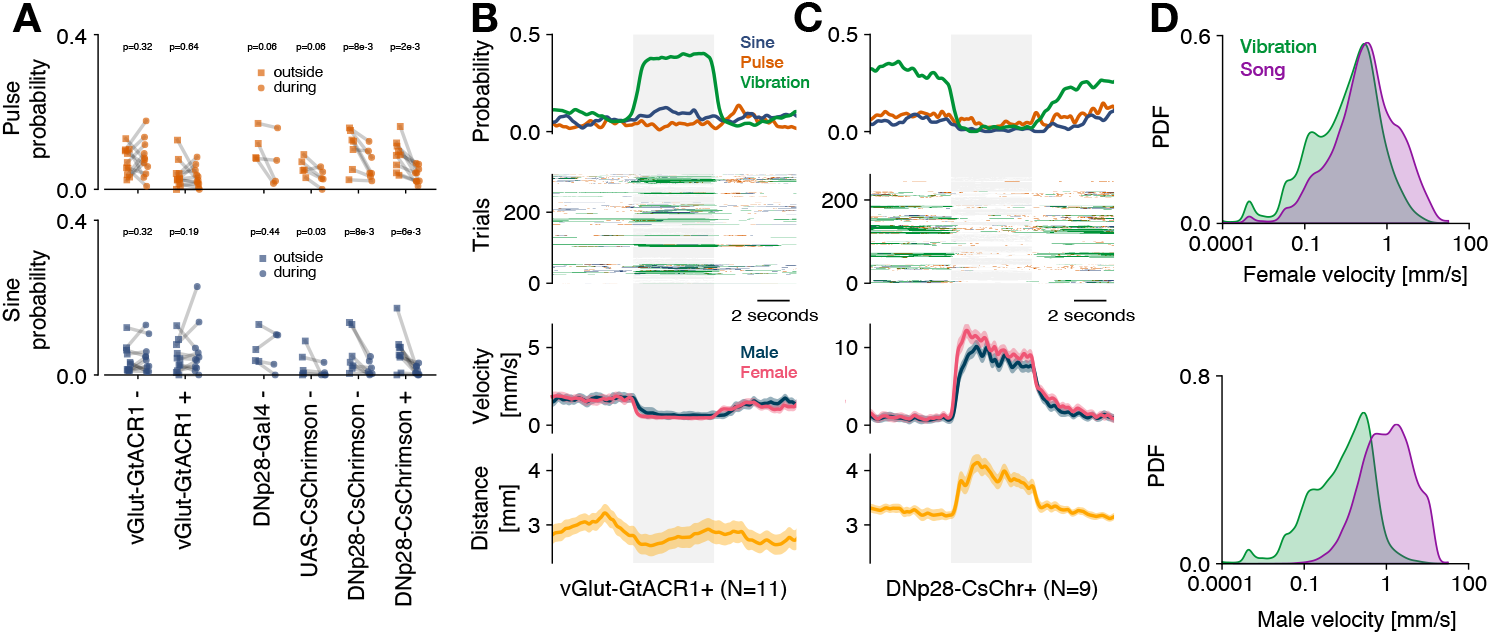
Manipulating female locomotion only weakly affects male singing. **A** Effect of female stopping (inactivation of all motor neurons with vGlut-GtACR1) and female acceleration (activation of DNp28 neurons with CsChrimson) on pulse song (top, orange), and sine song (bottom, blue). Same data as Fig. 3B, D but with sine and pulse song. Statistics compare the signal probabilities outside (squares) and during (circles) of optogenetic stimulation for each genotype. “+” and “-” after each genotype name indicate whether flies were fed all-trans retinal, a co-factor necessary for light sensitivity in Chrimson and GtACR1 that is present only in small amounts in regular food. P-values for vGlut-GtACR1 (+ and -) from a Wilcoxon test of the hypothesis that optogenetic stimulation increases signaling. P-values for the remaining genotypes from a Wilcoxon test of the hypothesis that optogenetic stimulation decreases signaling. **B, C** Trial-averaged probability of observing sine (blue), pulse (orange) and vibration (green) (top), single trial signaling (upper middle), male (blue) and female (pink) velocity (lower middle, line - mean, shaded area - standard error), and male-female distance (bottom, mean±standard error of the mean) during optogenetic inactivation of vGlut (B) and optogenetic activation of DNp28 (C). The time of optogenetic stimulation is marked as a grey shaded area. Inducing female stopping through vGlut inactivation drives vibration, but has no effect on distance and song (B). Inducing female acceleration suppresses vibrations and pulse and sine and increases the male-female distance. **D** Distributions of female (top) and male (bottom) velocity during song (purple) and vibration (green). Female velocities overlap more than male velocities, indicating that male movement determines the choice between song and vibration more than female movement.

**Figure S5:**
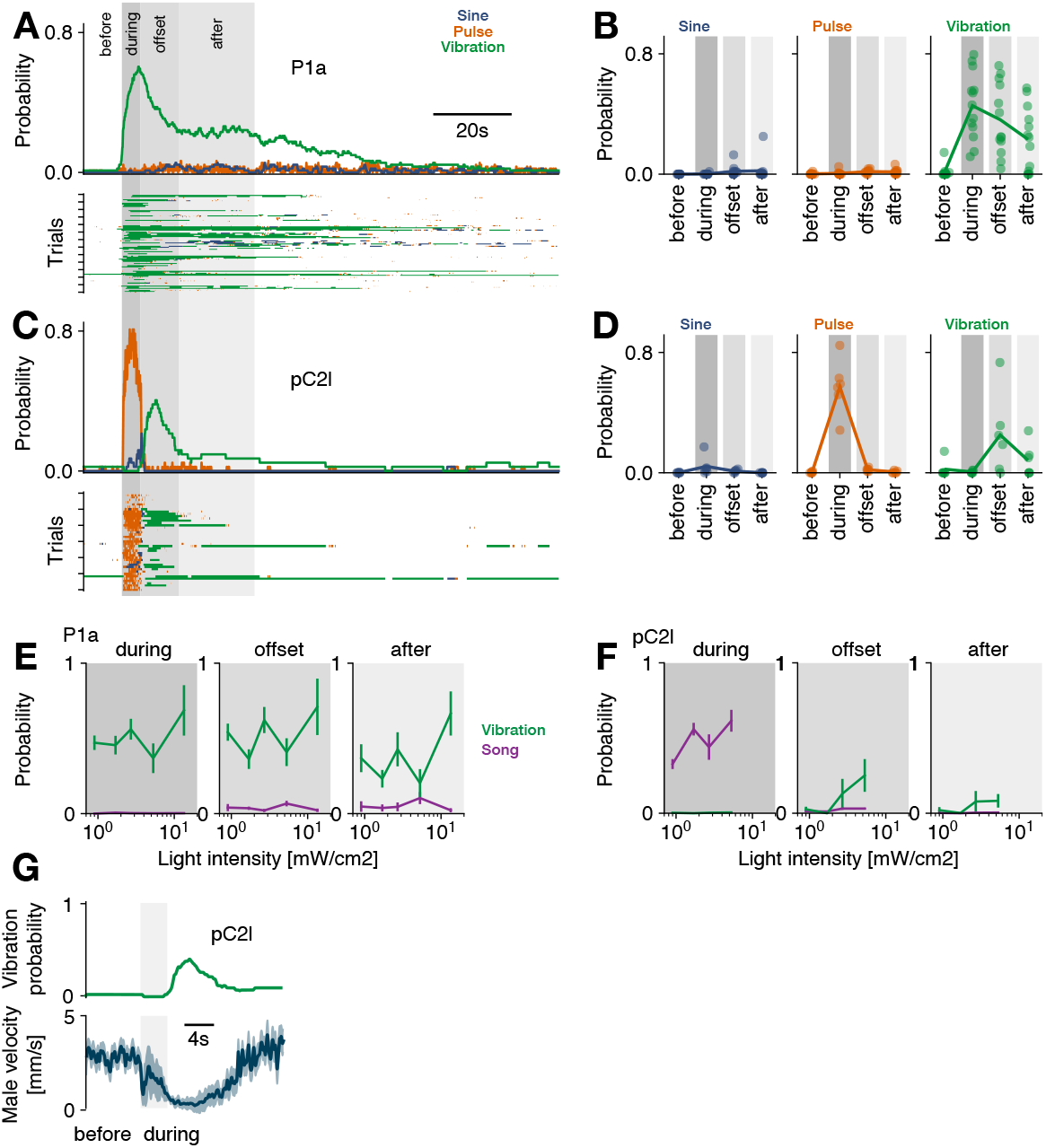
Activation of P1a and pC2l drives song and vibration. **A** Trial average probability (top) and single trial raster (bottom) for sine (blue), pulse (orange), and vibration (green) in response to optogenetic activation of P1a in solitary males (27 mW/cm^2^, N=13 flies, 7 trials/fly). Gray shaded areas delimit the epochs analysed in D. Same as Fig. 4C but song is split into sine and pulse modes. **B** Probability of observing sine (left), pulse (middle), and vibration (right) in different epochs surrounding P1a activation. **C** Same as A but for optogenetic activation of pC2l in solitary males (83 mW/cm^2^, N=6 flies, 7 trials/fly). **D** Same as B but for pC2l activation. **E, F** Probability of observing song (purple) and vibration (green) in different epochs surrounding the activation of P1a (E) or pC2l (F) at different intensities (625 nm). **G** Vibration probability (green) and male velocity (blue, mean±standard deviation over N=13 males with 7 trials each) in response to optogenetic activation of pC2l. Same data as C.

**Figure S6:**
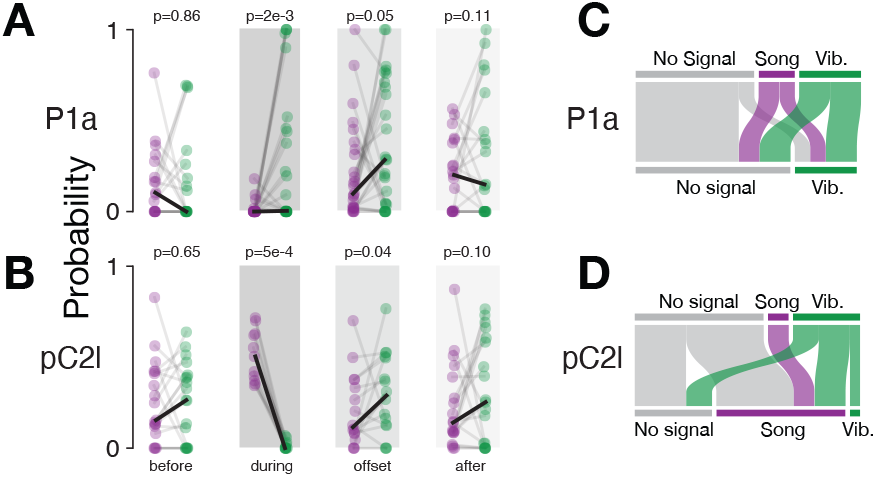
Activation of P1a and pC2l in males courting a female. **A, B** Comparison of the probability of song (purple) and vibration (green) upon activation of P1a (A) and pC2l (B) in a male courting a female. Same data as in Fig. 5C, D. P-values from Wilcoxon tests (before: two-sided; during: P1a more vibration than song, pC2l less vibration; offset and after: more vibration than song; all hypotheses based on the results of activation in solitary males in Fig. 4 C–F). **C, D** Transitions between song, vibration and silence when P1a (C) or pC2l (D) are activated optogenetically in males courting a female. After P1a activation, all males either vibrate or stop signaling. After pC2l activation, vibrating males tend to start singing or stop signaling.

**Figure S7:**
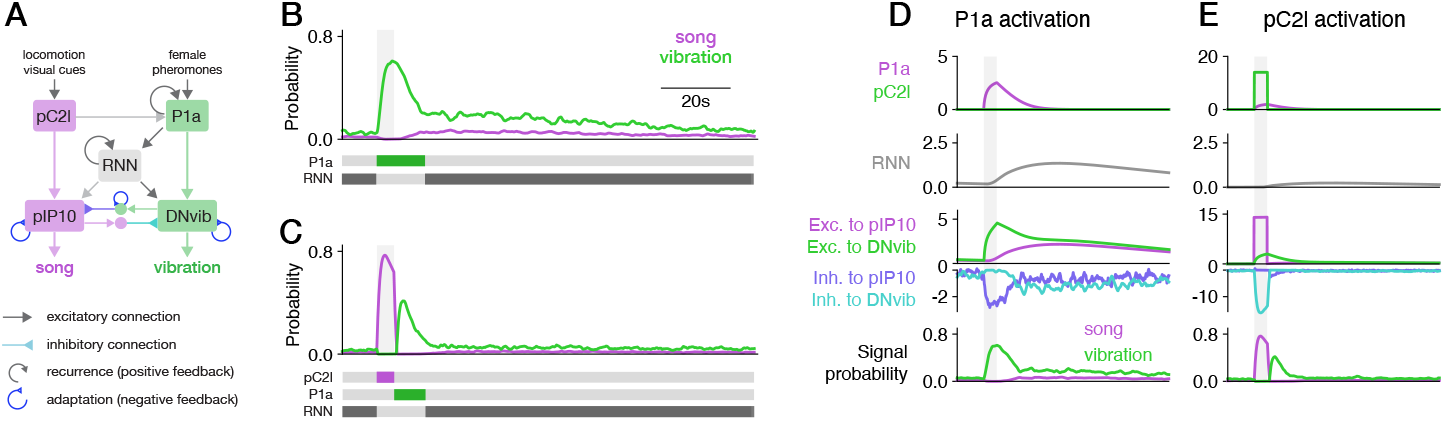
Detailed diagram of the network model. **A** Detailed network diagram for the model. Gray and blue arrows with straight and inverted heads indicate excitation and inhibition, respectively. Circular arrows indicate positive feedback (recurrence) and negative feedback (adaptation). Colors denote the signal driven during activation of each neuron (purple - song, green - vibration). **B, C** Schematic diagram of which neurons drive which signals in different phases during activation of P1a (B) and pC2l (C). P1a drives vibrations during and at the offset of P1a activation. pC2l drives song during activation. P1a drives vibration at the offset of pC2l activation. The recurrent neural network drives signaling in the persistent phase, starting 10 seconds after activation. **D** Activity of individual neurons in the model during activation of P1a. Optogenetic activation of P1a decays slowly because of intrinsic processes (purple, top) and induces persistent activity in the RNN (grey, 2nd row). DNvib is directly activated by P1a (green, 3rd row), which drives strong vibrations during and immediately after P1a activation (green bottom). The RNN kicks in later to provide persistent inputs to DNvib and to pIP10 (3rd row). Strong activation of the DNvib during P1a activation drives strong inhibition to pIP10 (violet, 4th row) and thereby suppresses song during P1a activation. Inhibition from pIP10 to DNvib only kicks in later (cyan, 4th row) and enables noise-induced switching between song and vibration during the persistent phase. **E** Optogenetic activation of pC2l drives pC2l activity but also weakly activates P1a (purple and green, top). The P1a activity is too weak to strongly activate the RNN (grey, 2nd row), thereby preventing persistent signaling. During pC2l activation, pIP10 is strongly activated by pC2l and drives singing (purple, 3rd row). At the same time pIP10 strongly inhibits DNvib (cyan, 4th row) which suppresses vibrations. DNvib gets input from the slower P1a activity, which outlasts the pC2l activity and the inhibition from pIP10 (green, 3rd row). The slowly decaying P1a activity then drives at the offset of pC2l activation (bottom).

**Figure S8:**
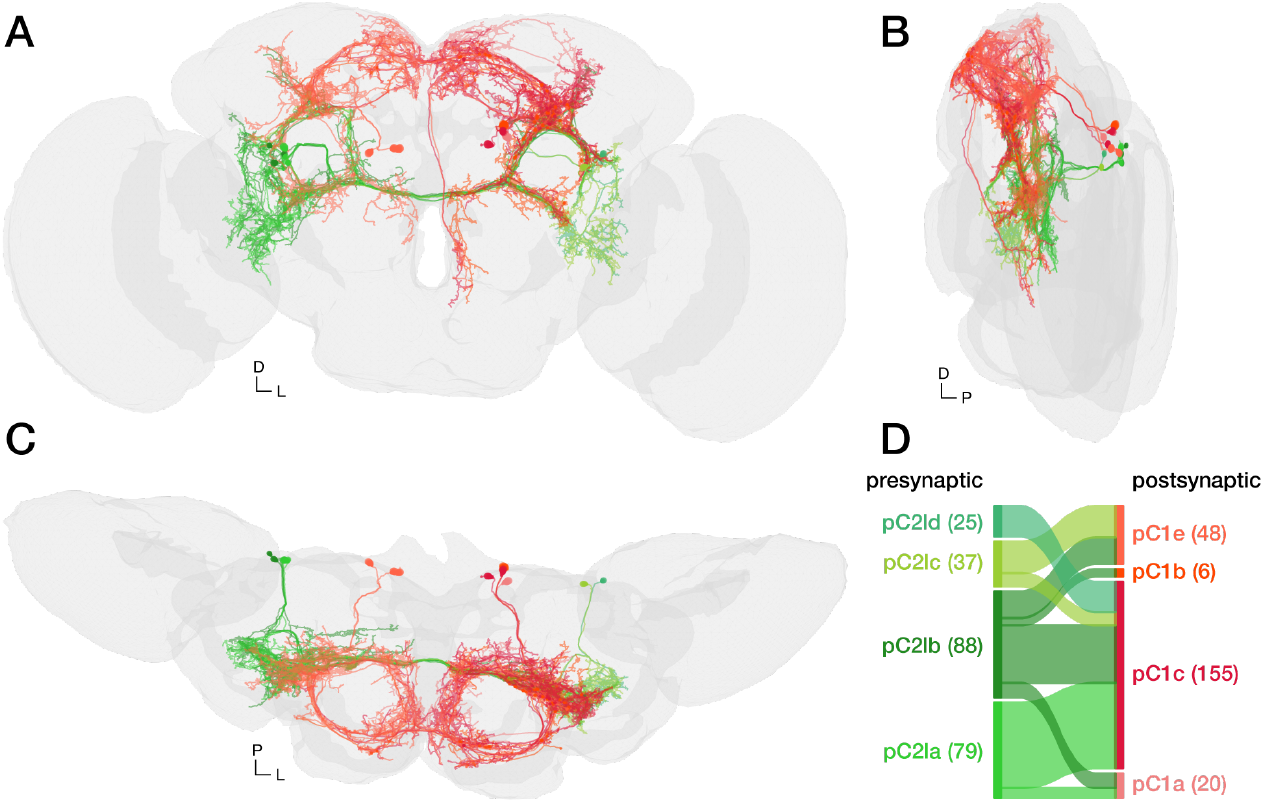
Connections between pC2l and pC1 in the flywire connectome. **A–C** Frontal (A), lateral (B), and dorsal (C) view of pC2l (green shades) connected to pC1 neurons (red shades) in the connectome of the female brain. The P1a neurons are a male-specific subtype of the pC1 neurons in the female. Different shades of green and red indicate different subtypes of pC2l (a–d) and pC1 (a–e), respectively (color code in D). Grey shows a volume rendering of the fly brain. **D** Connectivity between different subtypes of pC2l (presynaptic) and pC1 (postsynaptic) neurons. Line width is proportional to synapse count for each type of connection. Numbers beside each subtype indicate the number of outgoing (left) and incoming (right) synapses. In the female brain, there are in total 229 cholinergic synapses between 4 pC2l and 4 pC1 subtypes. It is thus likely that similar connections exist between pC2l and P1a in the male.

**Figure S9:**
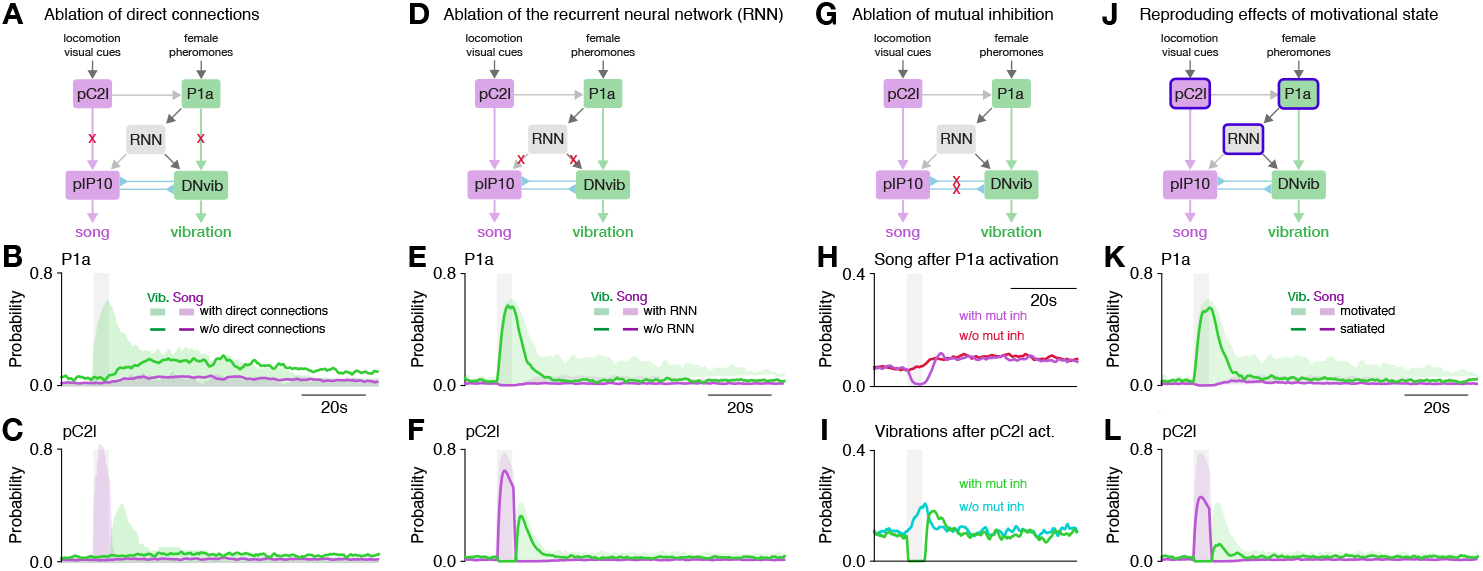
Ablation experiments and impact of motivation state in the circuit model. **A** Testing the role of direct connections between P1a and DNvib and between pC2l and pIP10 in the model through ablation (red crosses mark ablated connections). **B, C** Song (purple) and vibration (green) for activation of P1a (B) and pC2l (C) in an intact model (shaded areas) and in a model without direct connections to pIP10 and DNvib (lines) (compare data in Fig. 4C, E). Removing the direct connections removes the vibrations evoked during and shortly after activation of P1a (B) as well as the song and the vibration produced during and after pC2l activation (B). The sustained song and vibration are not affected by removal of the direct connections. Thus, the direct connections drive signals during and shortly after activation of pC2l and P1a. The latter effect arises from the slow decay of P1a activity. **D** Testing the role of the recurrent neural network (RNN) in the model by removing the connections from the RNN to pIP10 and DNvib (red crosses). **E, F** Song (purple) and vibration (green) for activation of P1a (E) and pC2l (F) in an intact model (shaded areas) and in a model without an RNN (lines) (compare data in Fig. 4C, E). Ablating the RNN strongly reduces the persistent signaling after activation in P1a but has otherwise only weak effects. Thus, the RNN drives signaling mainly during the persistent phase. **G** Testing the role of mutual inhibition in the network model by removing the inhibitory connections between pIP10 and DNvib (red crosses). **H, I** Song upon P1a activation (H) and vibrations upon pC2l activation (I) in an intact network (purple and green lines) and in a network without mutual inhibition (red and cyan lines) (compare data in Fig. 5C–D). Without mutual inhibition signals (song/vibration) are not suppressed during activation of P1a/pC2l. **J** Modeling the impact of sexual satiation on the circuit. Sexual satiation was modeled by reducing the excitability in pC2l, the slow decay P1a as well as the recurrent excitation in the RNN. **K, L** Song (purple) and vibration (green) for activation of P1a (K) and pC2l (L) in naive, sexually motivated males (shaded areas) and in sexually satiated males (lines). In the model responses of pC2l to activation are reduced, as are the persistent vibrations after activation of P1a and pC2l. This is consistent with the experimental data in Fig. 5I–J.

